# Metabolic and transcriptomic analyses of nectaries reveal differences in the mechanism of nectar production between monocots (*Ananas comosus*) and dicots (*Nicotiana tabacum*)

**DOI:** 10.1101/2024.06.21.597291

**Authors:** Thomas Göttlinger, Marcello Pirritano, Martin Simon, Janina Fuß, Gertrud Lohaus

## Abstract

**Background:** Nectar is offered by numerous flowering plants to attract pollinators. To date, the production and secretion of nectar have been analyzed mainly in eudicots, particularly rosids such as *Arabidopsis*. However, due to the enormous diversity of flowering plants, further research on other plant species, especially monocots, is needed. *Ananas comosus* (monocot) is an economically important species and ideal for such analyses because it produce easily accessible nectar in sufficient quantities. In addition, the analyses were also carried out with *Nicotiana tabacum* (dicot, asterids) for comparison.

**Results:** We performed transcriptome sequencing (RNA-Seq) analyses of the nectaries of *Ananas comosus* and *Nicotiana tabacum*, to test whether mechanisms described for nectar production and secretion in *Arabidopsis* are also present in these plant species. The focus of these analyses is on carbohydrate metabolism and transport (e.g., sucrose-phosphate synthases, invertases, sucrose synthases, SWEETs and further sugar transporters). In addition, the metabolites were analyzed in the nectar, nectaries and leaves of both plant species to address the question whether concentration gradients for different metabolites exist between the nectaries and nectar.

The nectar of *N. tabacum* contains large amounts of glucose, fructose and sucrose, and the sucrose concentration in the nectar appears to be similar to the sucrose concentration in the nectaries. Nectar production and secretion in this species closely resembles corresponding processes in some other dicots, including sucrose synthesis in nectaries and sucrose secretion by SWEET9.

The nectar of *A. comosus* also contains large amounts of glucose, fructose and sucrose and in this species the sucrose concentration in the nectar appears to be higher than the sucrose concentration in the nectaries. Furthermore, orthologs of SWEET9 appear to be generally absent in *A. comosus* and other monocots. Therefore, sucrose export by SWEETs from the nectaries into the nectar can be excluded, rather, other mechanisms, such as active sugar export or exocytosis, are more likely.

**Conclusion:** The mechanisms of nectar production and secretion in *N. tabacum* appear to be largely similar to those in other dicots, whereas in the monocotyledonous species *A. comosus*, different synthesis and transport processes are involved.

## Background

Floral nectar contains several compounds, mainly sugars such as glucose, fructose and sucrose but also, smaller amounts of amino acids, inorganic acids and secondary compounds [1–3]. The nectar composition varies greatly depending of the plant species and is often related to the type of pollinator [4–6]. Floral nectar is produced by specialized glands called nectaries [7]. To date, different models for nectar production and secretion, such as the eccrine or the granulocrine model have been proposed [8, 9].

For some plant species, including *Arabidopsis*, an eccrine secretion mode has been described, wherein various transport proteins and enzymes are involved in nectar production [10, 11]. Three types of sugar transporters could be involved in sugar uptake into and export from the nectaries. Sugar transporter proteins or hexose transporters (STPs/HTs) are H^+^/monosaccharide symporters and these proteins are commonly found in plant sink organs, where they are located in the plasma membrane [12]. Sucrose uptake transporters (SUTs) are H^+^/sucrose symporters that are categorized into four or five types based on sequence homology and biochemical properties [13, 14]. Most SUTs are found in the plasma membrane, but SUT4-type proteins are also localized in the tonoplast [14]. Sugar will eventually be exported transporters (SWEETs) are uniporters divided into four clades, with clades I and II mainly transporting hexoses and clade III mainly transporting sucrose, while clade IV contains tonoplast-localized hexose transporters [15, 16].

Because most nectaries are non-photosynthetic sink tissues, they rely on phloem-derived sugars from source tissues, such as leaves [17, 18]. Sucrose is either symplasmically transported from the phloem through plasmodesmata into the nectary parenchyma cells or can be transported into the apoplasm. In cucumber, *CsSWEET7a* is highly expressed in the receptacle and nectary tissues of flowers just before anthesis and it may play a role in phloem unloading of sucrose into the apoplasm [19]. From the apoplasm, sucrose could be actively taken up into the nectaries through the activity of sugar transporters such as SUTs or after hydrolysis by cell wall invertases by STPs/HTs [18].

In various plant species, starch accumulates in the parenchyma cells of the nectaries in the early stages of nectary maturation [20–22]. During nectar secretion, starch is converted into sugars [20, 23, 24] and various enzymes are involved in this process, e.g. sucrose phosphate synthase (SPS; E.C. 2.4.1.14) [11]. In addition to nectar sugars derived from starch degradation in nectaries, phloem-derived sucrose can used directly for nectar production without prior storage as starch, as shown for tobacco [20] or squash [22].

The phloem sap of many plant species contains mainly sucrose, while hexoses are typically present only in very small concentrations [17, 25]. The proportion of hexoses in nectar therefore depends on sucrose-cleaving enzymes in the nectaries or during secretion. In plants, sucrose synthases (SUS; glycosyltransferase; EC 2.4.1.13) catalyze the reversible conversion of sucrose and UDP into fructose and UDP-glucose. Whether sucrose degradation or sucrose synthesis is promoted *in vivo* depends on the concentrations of the substrates and products [26]. Sucrose synthases exist in different isoforms with various biochemical properties and they have been observed either in the cytosol or associated with the plasma membrane [26]. Invertases (*β*-fructofuranosidases; EC 3.2.1.26) catalyze the irreversible sucrose hydrolysis into glucose and fructose. Invertases can be classified into three groups: neutral/alkaline invertases (NINVs), vacuolar invertases (VINVs), and extracellular invertases which are bound to the cell wall (CWINVs) [27, 28]. Sucrose hydrolysis during nectar secretion by CWINV4 has been shown in *Arabidopsis* and other plant species [11, 29].

For the export of sucrose from nectaries into nectar, the plasma membrane-localized sucrose transporter SWEET9 is essential, e.g. in *Arabidopsis* [11]. SWEET9 functions as a bidirectional and facilitated diffusion transporter for sucrose and its activity is dependent on the sucrose concentration gradient [11]. Therefore, the export of sucrose is only possible if the cytosol of the parenchyma cells of the nectaries contains higher sucrose concentrations than the nectar. The question still remains whether AtSWEET9 is able to secrete such high amounts of sugar and whether other mechanisms are also involved in this process [30]. In plant species with hexose dominant nectar, e.g. in Brassicaceae such as *Arabidopsis* or *Brassica*, the extracellular hydrolysis of sucrose into hexoses by cell wall invertases (CWINV4) could be the driving force for sucrose efflux [11, 29]. However, this does not apply to species that secrete nectar with high sucrose concentrations; they require alternative mechanisms for sucrose efflux. This may include the production of very high concentrations of sucrose in the nectaries, similar to the concentration in the nectar, due to induced sucrose synthesis during secretion and/or a reduced sucrose degradation [22]. In addition, the sucrose concentration may be very high in certain parts of the nectaries, especially in cells that secrete nectar.

Extracellular sucrose hydrolysis also increases the osmotic gradient that allows the secretion of water from nectaries into nectar [11, 29]. It is not yet known whether plant aquaporins, known as Plasma membrane Intrinsic Proteins (PIPs) [31], which enable rapid movement of water molecules across cell membranes, are also involved in water secretion in nectaries [9].

The granulocrine secretion mode is also discussed for some plant species, in which sugar secretion is mediated by exocytosis [7, 32]. According to this model, metabolites are packaged into vesicles in the outer cells of nectaries by dictyosomes or the endoplasmic reticulum (ER). The vesicles then fuse with the plasma membrane and release the metabolites to nectar [7]. SNARE-domain containing proteins are characterized by a particular SNARE motif (soluble *N*-ethylmaleimide-sensitive factor adaptor protein receptors). They play an important role in vesicle-associated membrane fusion events in transport processes, including exocytosis [33]. Monocots and dicots encode a high number of SNARE proteins, which are divided into different classes: Qa, Qb, Qc, and Qb+Qc, which are t-SNAREs (target membrane-associated SNAREs), and R-type SNAREs, which are v-SNAREs (vesicle-associated SNAREs) [34]. One could hypothesize that SNAREs might be involved in the secretion of nectar sugars by exocytosis, but experimental evidence of this is still lacking.

Most experiments on nectar production and secretion have been carried out with eudicots, particularly *Arabidopsis*. However, due to the enormous diversity of flowering plants, further research on other plant species, especially monocots, is required [9]. Therefore, to compare the mechanism underlying nectar production and secretion in monocots and dicots, pineapple (*Ananas comosus* L. Merr.; Bromeliaceae) and tobacco (*Nicotiana tabacum* L., Solanaceae) were chosen for the experiments. Both species are economically important and produce easily accessible nectar in sufficient quantities. *A. comosus* is a diploid species with a relatively small genome size of 526 Mb [35], whereas *N. tabacum* is an allotetraploid species with a large 4.5 Gb genome containing a high proportion of repetitive elements [36]. Furthermore, *N. tabacum* uses the C3 photosynthetic pathway and *A. comosus* is a CAM plant.

The type of nectaries also differs between both plant species. In *A. comosus* and other bromeliads, floral nectar is produced by septal nectaries which are located in the basal part of the ovary and are formed by incomplete fusion of the carpels [37]. In *Ananas ananassoides*, the septal nectaries are not vascularized, but they are connected to numerous vascular bundles in the ovaries [32]. In *N. tabacum* and other Solanaceae the floral nectar is produced by gynoecial nectaries which are located on the basal side of the gynoecium [38].

To gain further insight into the molecular mechanisms involved in nectar production and secretion by *Nicotiana tabacum* and *Ananas comosus*, RNA-Seq analyses were performed to examine the nectary transcriptome. The focus of these analyses was on carbohydrate metabolism and transport. For comparison, RNA-Seq analyses were also carried out on the leaves of both plant species and furthermore, the metabolites in the nectar, nectaries and leaves of both plant species were analyzed to address the question of whether concentration gradients for different metabolites exist between the nectar and nectaries. During nectar production, the cleavage of sucrose to hexoses is necessary; therefore, the activities of the involved enzymes were also determined.

## Methods

### Plant material

*A. N. tabacum* seeds were obtained from NiCoTa (Rheinstetten, Germany). Each plant was potted in a single 5 L pot with compost soil and grown in a greenhouse at the University of Wuppertal (Germany). Cultivation was carried out with a 14-h-light/10-h-dark cycle, an irradiance of approximately 300 µmol photons m^-2^ s^-1^ and a temperature regime of 25 °C day/18 °C night.

*Ananas comosus* plants grown in tropical glasshouses in the Zoological-Botanical Garden Stuttgart (Germany) and at the University of Wuppertal (Germany) were used for the different analyses. The plants were grown in Brill Pro Verde substrate (Georgsdorf, Germany) enriched with 30 % pine bark under the same cultivation conditions as those used for the tobacco plants.

### Collection of leaf tissue, nectaries and nectar

For each tissue (leaf, nectary, nectar) at least three samples of *Ananas comosus* and *Nicotiana tabacum* were collected. All samples of plant material were harvested 3-4 h after anthesis. All the samples were immediately frozen in liquid nitrogen and stored at -80 °C until further analysis.

For the leaf material, samples (∼200 mg) were taken from leaves with a razor blade.

Each sample (∼100 mg) of nectary tissue comprised 20 to 30 nectaries, depending on the species. To collect the nectaries from *Ananas comosus*, the gynoecia were extracted from the flowers, and the septal nectary tissue was dissected with a scalpel and rinsed with ultrapure water to remove external sugars [39, 40]. The nectary tissue of *Nicotiana tabacum* was dissected with a scalpel from the flower at the base of the ovary, as this is recognizable by its orange coloring caused by β-carotene. Afterwards, the tissue was also washed with ultrapure water to remove external sugars [41].

After anthesis, each nectar sample was collected from a single flower using a micropipette [6]. The volume of nectar from the flowers varied between 10 and 50 µl. To avoid possible pollen contamination, the nectar samples were examined microscopically. Furthermore, microbial contamination was evaluated according to an assay for microbial contamination [5]. The test revealed no microbial contamination in the nectar samples from either species.

### Water content of the nectaries and leaves

Leaves and nectaries were weighed, dried and reweighed to determine the water content in these tissues. The water content was calculated from the ratio between the dry weight and the fresh weight [41].

### Extraction of soluble metabolites from leaf and nectary tissue

Chloroform-methanol-water extraction has been used to extract soluble metabolites, such as sugars and amino acids, from nectaries or leaves [42]. For this purpose, 200 mg of milled leaf material and 100 mg of milled nectary material frozen in liquid nitrogen were used.

### Analysis of sugars and free amino acids in nectar, nectary, and leaf tissue

The collected nectar samples, extracted nectaries, and extracted leaf tissue were analyzed by using HPLC to determine the concentration and composition of sugars and amino acids.

The concentrations of the different sugars in the plant materials were determined via an ICS-5000 HPIC system (Thermo Fisher Scientific). For the analysis, the sugars were eluted isocratically using an anion exchange column and a pulse amperometric detector for data collection [17].

An Ultimate 3000 HPLC system (Thermo Fisher Scientific) was used for the detection of amino acids. After separation on a reversed-phase column (Merck LiChroCART® 125-4 using Superspher® 100 RP-18 endcapped), free amino acids (alanine, arginine, aspartate, asparagine, glutamate, glutamine, glycine, histidine, isoleucine, leucine, lysine, methionine, phenylalanine, proline, serine, threonine, tryptophan, tyrosine, valine) in the different plant materials were analyzed by a fluorescence detector.

By using a calibration curve for each component, the chromatograms were evaluated by an integration program (Chromeleon 7.2). By measuring the sugar content in the leaves and nectaries in µmol g^-1^ fresh weight (FW) and the water content of the leaves and nectaries, it was also possible to determine the sugar concentration (mM) in both tissues [6, 41].

### Analyses of starch in leaves and nectaries

The insoluble residues of the chloroform-methanol-water extraction of leaf and nectary tissue samples as described above were treated with KOH, α-amylase and amyloglucosidase to cleave the starch into glucose [43]. Aliquots (50 µl) each incubation mixture were analyzed spectrophotometrically for glucose [43]. The starch content was calculated as milligrams of glucose equivalent per gram of fresh weight.

### Enzyme assays for cell wall invertase (CWINV), vacuolar invertase (VINV) and neutral invertase (NINV)

To measure the enzyme activity of the three different invertases, a protein extraction was first carried out on 50 mg of nectary material [41]. The invertase reaction of CWINV was conducted with an aliquot of insoluble protein extract added to 0.6 M sucrose and 0.125 M sodium acetate (pH 5.0). For vacuolar and neutral invertases, an aliquot of soluble protein extract was added to 0.6 M sucrose and 0.125 M sodium acetate, pH 5.0 (VINV) or pH 7.5 (NINV). To stop the enzyme reaction the solution was boiled for 10 min. Afterwards, the amount of glucose released during each reaction was quantified optically by the reaction of hexokinase and glucose-6-phosphate dehydrogenase [44]. The activity of the enzymes was calculated by the activity in the sample minus the activity in the blank. Blanks in which the reaction mixture was immediately inactivated by heat without incubation were used.

### Analysis of sucrose synthase (SUS) enzyme activity

The cleavage of sucrose was investigated to determine the activity of sucrose synthases (SUSs). The soluble and insoluble proteins were extracted from 50 mg of fine-milled nectary tissue [45]. The extracted protein fraction was added to 100 mM sucrose, 4 mM UDP, and 20 mM HEPES (pH 7.0) to incubate this solution for the cleavage of sucrose. After 40 min of incubation at room temperature, the enzymatic reaction was stopped by boiling the solution for 10 min. The products of the incubation were determined by coupled optical enzymatic assays [46]. The activity of the enzyme was calculated by the activity in the sample minus the activity in the blank. Blanks in which the reaction mixture was immediately inactivated by heat without incubation were used.

### RNA isolation, library preparation and sequencing

Approximately 50 mg of nectary tissue and 200 mg of leaf tissue were milled to a fine powder in liquid nitrogen. Three tissue samples each of *A. comosus* and *N. tabacum* were extracted using a modified protocol [47]. Denaturing agarose-gel electrophoreses was used to assess the integrity of the extracted RNA. Furthermore, the purity and concentration were verified by a UV‒vis spectrophotometer at 260 and 280 nm (NanoDrop™ One, Thermo Fisher Scientific). DNase I digestion and RNA cleaning were performed with an RNA Clean & Concentrator kit (Zymo Research Europe). The concentration and integrity of the RNA were verified using a Qubit^TM^ RNA HS Assay Kit (Invitrogen) and denaturing agarose-gel electrophoreses, respectively. Afterwards, poly-A enrichment was carried out with 1 µg of this RNA using an mRNA isolation method (NEBNext^®^ Poly(A) mRNA Magnetic Isolation Module). Library preparation was performed using the NEBNext^®^ Ultra^TM^ II DNA Library Prep Kit for Illumina^®^ using 11 PCR cycles according to the manufacturer’s instructions. A Qsep1 Bio-Fragment Analyzer (BiOptic Inc.) with a Standard DNA Cartridge Kit was used to check the library for adaptor contamination and size distribution. The prepared libraries were multiplex sequenced on an Illumina NovaSeq 6000 platform in paired-end mode. Reads were demultiplexed and the adapter and quality trimming were carried out with the software tool Trimgalore (version 0.6.5, www.bioinformatics.babraham.ac.uk), which uses the tool Cutadapt Wrapper (version 1.18, www.cutadapt.readthedocs.io) [48].

### Analysis and visualization of transcriptome data

The trimmed sequences of *A. comosus* and *N. tabacum* were paired and subsequently mapped with the memory-efficient tool Bowtie2 to the corresponding genome using Geneious Prime^®^ software (version 2024.0.4, www.geneious.com) [49]. For the pineapple sequences, the whole-genome shotgun sequence of Ming et al., 2015 [50], and for the tobacco sequences, the genome sequence of Edwards et al., 2017 [51], were used. The mapped reads were used to calculate normalized expression levels for RNA from each sample and tissue. This allows the TPM (transcript per million) to be determined for each gene as a measure of the level of transcription. These values were used to compare the genes within a gene group. The number of mapped reads for a gene depends on the expression level, the gene length and the sequencing depth. To compare the nectary and leaf tissue of each species, DESeq2 (1.42.1) in Geneious Prime^®^ software was used to normalize the dependencies of these and calculate the differential expression between the two samples [52]. As a result of this calculation, the fold change (Fc) in expression between the sample condition A (leaf) and sample condition B (nectary) was expressed as a log2 ratio. In this way it is possible to compare the expression levels of a gene set between different tissues of a species. The genes of each species had to meet the criteria: (1) log2 ratio > 1 or log2 ratio < -1 to be considered differentially expressed between the compared tissues. Differential expression was considered to be statistically significant if the adjusted *p* value was less than 0.05. To visualize the differentially expressed gene loci, a volcano diagram was created with the fold change plotted against the absolute confidence. In addition, Principal Component Analysis (PCA) was performed to visualize the variation between the gene expression levels of samples and to assess the quality of the experimental design. Furthermore, the transcript levels (TPMs) of the different genes involved in nectar production and secretion in each sample (leaf and nectary) were used to plot an expression heatmap of *N. tabacum* and *A. comosus* to visualize the variation between samples and genes. An expression heatmap was generated with the R software (version 4.4.0, www.r-project.org). The TPM data used from the leaf and nectary samples were normalized for each species using the ‘scale()’ function of R software, where each element of the data matrix was scaled by subtracting the mean of the matrix and dividing by the standard deviation.

### Gene Ontology (GO) enrichment analysis

Since the two genomes used for *A. comosus* and *N. tabacum* did not have comprehensive annotations, the eggNOG-mapper tool (www.eggnog-mapper.embl.de) was used for rapid functional annotation [53, 54]. The application of this tool added the corresponding gene ontology (GO) terms to the genes, providing information on the function of the genes. These data were used as background GO data for the GO enrichment analysis. In general, the analysis revealed GO terms that were over- or underrepresented in a gene set. These terms are grouped into three categories: molecular function, cellular component, and biological process. Therefore, for each species, genes with a log2 ratio greater than 1 and less than -1 were used separately in the analysis. The bioinformatics platform tool TBtools-II was used to perform a GO enrichment analysis [55]. As a result, each gene set was visualized by plotting the absolute confidence (- log10 adjusted *p* value) of the GO terms in relation to three categories.

### Phylogenetic analysis

The software Mega (version 11.0.13, www.megasoftware.net) [56] with the maximum likelihood method and 1,000 bootstrap iterations was used to construct the phylogenetic trees for the respective gene groups. The accession numbers used to identify the genes from *Arabidopsis thaliana*, *Nicotiana tabacum*, *Oryza sativa*, and *Ananas comosus* in the NCBI database can be found in Supplementary Table S2-5.

### Statistical analysis

The significance between metabolite and starch concentrations and enzyme activities, was determined in two groups by using *t*-tests, and in more than two groups using one-way ANOVA, followed by Tukey *post-hoc* tests. Statistical analyses were performed with the R software (version 4.4.0, www.r-project.org).

## Results

### Sugar, starch and amino acid concentrations in leaves, nectaries, and nectar

*Nicotiana tabacum* and *Ananas comosus* are day-flowering species and nectar, nectaries and leaves were collected from plants of both species 3-4 hours after anthesis. The concentrations of sugars, starch and amino acids in the nectar, nectaries, and leaves of *N. tabacum* and *A. comosus* are shown in Figure 1. In both plant species, the leaves, nectaries, and nectar contained mainly of glucose, fructose and sucrose. The sugar concentrations in the nectar were determined in millimolar (millimole per liter). To allow easier comparisons between nectar and nectaries, the sugar concentrations in the nectaries were also calculated (in millimolar concentrations) by measuring the sugar content in the nectaries in micromoles per gram fresh weight and the water content of the nectaries (Supplementary Table S1). Sugar concentrations in leaves were also calculated in this way. The concentration of the sum of sugars was highest in nectar, followed by nectaries, and lowest in leaves (Fig. 1A, B). The mean sugar concentration for both species was about five- to nine-fold higher in the nectaries (310, 228 mM) than in leaves (57, 25 mM) and about three- to five-fold higher in the nectar (907, 1063 mM) than in the nectaries.

**Figure 1:**
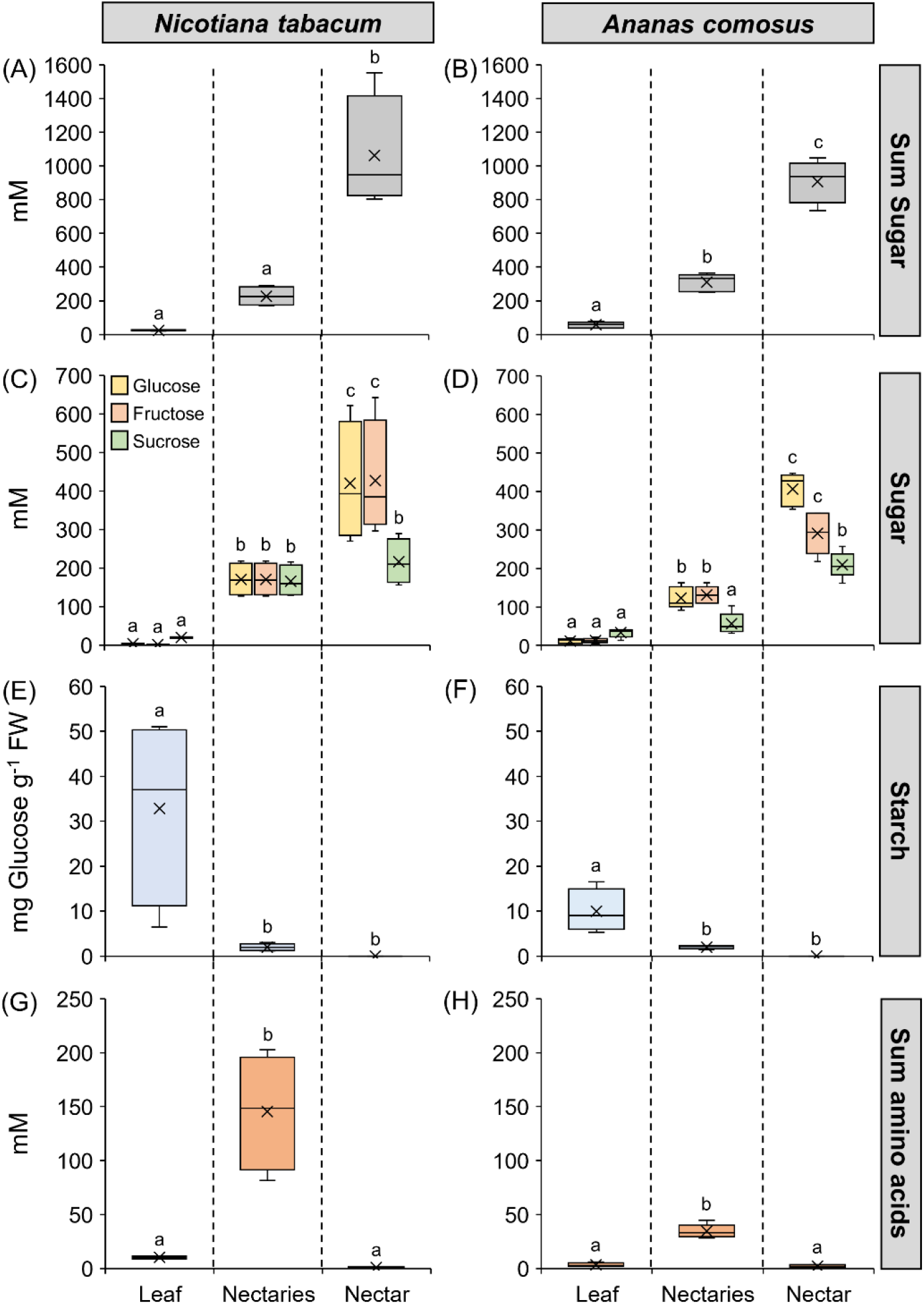
Composition of sugars, starch, and sum of amino acids in leaves, nectaries, and nectar. The metabolites are separated in the boxplot diagram (A-H) by the three tissues (leaf, nectary, nectar), and between *Nicotiana tabacum* and *Ananas comosus*. The shown data include four samples for each tissue of each species (n = 4). Different letters represent significant differences in metabolites between leaves, nectaries, and nectar (Tukey’s HSD; *p* < 0.05).

The leaves of *N. tabacum* contained more sucrose than hexoses, and the sucrose-to-hexoses ratio (4.2) was approximately nine-fold higher than that in the nectaries (0.49). In nectar, the ratio was lower (0.26) than that in nectaries, meaning that nectar contained more hexoses than sucrose compared to nectaries (Table 1; Fig. 1C). In *A. comosus*, the sucrose-to-hexoses ratio was approximately five-fold higher in leaves (1.4) than in nectaries (0.22), whereas in nectar, the ratio was slightly higher (0.30) than in nectaries. This means that nectar contained more sucrose than hexoses compared to nectaries (Table 1; Fig. 1D).

**Table 1:** Sugar ratios in leaves, nectaries and nectar (calc. from mM) The data were derived from Fig. 1, and the values of all the measuring points were averaged.

| Species | Leaf-sugar-ratio<br>[S/(G+F)] | Nectary-sugar-ratio<br>[S/(G+F)] | Nectar-sugar-ratio<br>[S/(G+F)] |
| --- | --- | --- | --- |
| *N. tabacum* | 4.2 | 0.49 | 0.26 |
| *A. comosus* | 1.4 | 0.22 | 0.30 |

In *N. tabacum*, the starch content in nectaries was 2.0 ± 0.8 mg g^-1^ FW and in *A. comosus* 2.0 ± 0.4 mg g^-1^ FW (measured as glucose equivalent; Fig. 1E, F). The starch content was much higher in the leaves of both species (10-33 mg g^-1^ FW).

The amino acid concentrations in leaves and nectaries were derived analogously to the sugar concentrations from the amino acid contents in micromole per gram fresh weight and the corresponding water contents (Supplementary Table S1). The highest amino acid concentration was found in nectaries of both species (Fig. 1G, H), with the concentration in the nectaries of *N. tabacum* (156 ± 56 mM) being much higher than in *A. comosus* (30.1 ± 6.3 mM). In the leaves of both species, the amino acid concentration was in the lower millimolar range (3.3, 1.0 mM), and the lowest concentration was found in nectar (1.5, 1.0 mM).

### Activity of invertase (INV) and sucrose synthase (SUS) in the nectaries

The activities of the sucrose cleavage enzymes invertase and sucrose synthase were determined in the nectaries of both species. The measured cell wall invertase activity in nectaries was 31.6 ± 9.0 U g^-1^ FW in *N. tabacum* and 4.7 ± 0.3 U g^-1^ FW in *A. comosus* (Table 2). This means that the activity in *N. tabacum* was seven-fold higher than that in *A. comosus*. Soluble acid invertases and soluble neutral invertases were also active in the nectaries, but the measured activities were lower (less than 2 U g^-1^ FW each) and no differences were found between *N. tabacum* and *A. comosus* (Table 2). In the nectaries of *N. tabacum* the sucrose synthase (SUS) activity was 3.8 ± 0.7 U g^-1^ FW, whereas a four-fold-lower activity of 0.9 ± 0.1 U g^-1^ FW was measured in the nectaries of *A. comosus* (Table 2).

**Table 2:**
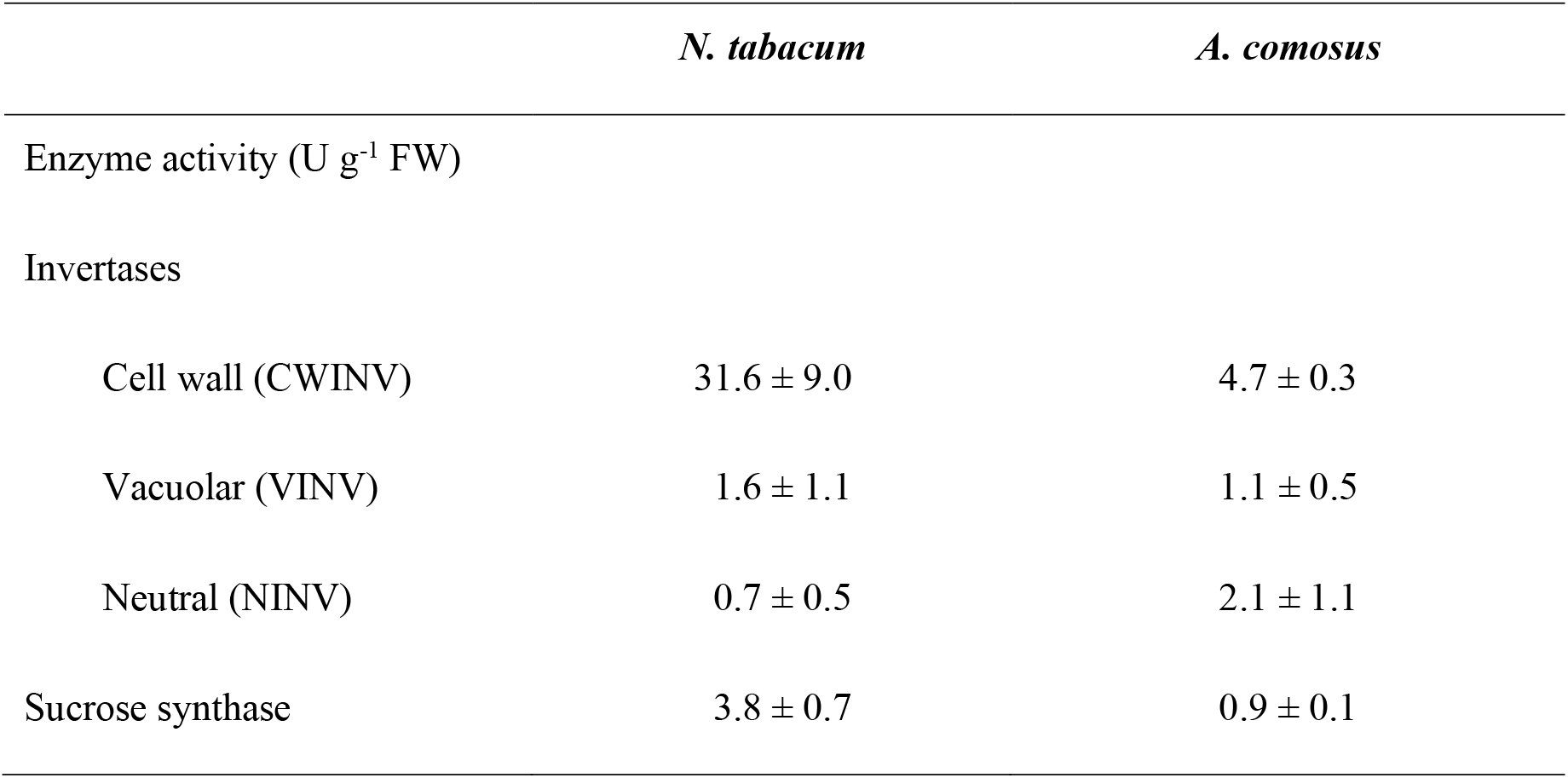
Activities of sucrose-cleavage enzymes in the nectaries of *N. tabacum* and *A. comosus*. The results are the means ± SDs of at least three samples from different plants.

|  | *N. tabacum* | *A. comosus* |
| --- | --- | --- |
| Enzyme activity (U g<sup>-1</sup> FW) |  |  |
| Invertases |  |  |
| Cell wall (CWINV) | 31.6 ± 9.0 | 4.7 ± 0.3 |
| Vacuolar (VINV) | 1.6 ± 1.1 | 1.1 ± 0.5 |
| Neutral (NINV) | 0.7 ± 0.5 | 2.1 ± 1.1 |
| Sucrose synthase | 3.8 ± 0.7 | 0.9 ± 0.1 |

### Overall gene expression profiles in the nectaries and leaves of *N. tabacum* and *A. comosus*

To link different genes to nectar production and secretion, the gene expression profiles of the nectaries as opposed to the leaves of *A. comosus* and *N. tabacum* were collected via RNA-Seq analyses. Comparison of the individual expression levels revealed in a log2 ratio (fold change) for 87% of the genes in the *N. tabacum* genome and 97% in the *A. comosus* genome. DESeq2 analysis was conducted to identify the most highly differentially expressed gene (DEG) loci, which were visualized in a volcano plot for each species (Supplementary Fig. S1A, D). To count as differentially expressed, genes of each species had to meet two criteria to be defined as DEGs: (1) log2 ratio > 1 (which are genes showing higher expression in nectaries than leaves) or log2 ratio < -1 (which are genes showing higher expression in leaves than nectaries) and (2) adjusted *p* value < 0.05. This eliminates variation at low expression levels, but also eliminates outliers at high expression levels. The orange dots in the volcano plots represent genes that fulfil this requirement, whereas the black dots represent genes that do not fulfil at least one requirement (Supplementary Fig. S1A, D). In general, both volcano plots show a large number of DEGs. If the log2 ratio (fold change) is positive, the DEG is up-regulated in the nectary tissue, meaning that the genes are more highly expressed in nectaries than leaves. Conversely, a negative log2 ratio indicates that the DEGs are down-regulated in the nectary tissue, meaning that the genes are more highly expressed in leaves than nectaries. In *N. tabacum*, more DEGs are down-regulated (6491 genes) than up-regulated (4978 genes) in the nectaries compared to the leaves. According to the volcano plot of *A. comosus*, more DEGs appeared to be up-regulated (5924 genes) than down-regulated (4365 genes) in the nectaries compared to the leaves (Supplementary Fig. S1C, F). Principal component analysis (PCA) also revealed that the nectar and leaf samples of both species were very different from each other while showing minimal variation between replicates at the same time (Supplementary Fig. S1B, E). For both species, the first principal component (PC1) accounted for 99% of the variation, while the second principal component (PC2) accounted for less than 1%. To further test the reliability of the RNA-Seq data and the DEGs, the expression levels of the transcription factor gene CRABS CLAW (*CRC*) were examined. *CRC* belongs to the YABBY family and its expression is associated with the development of carpels and nectaries [57]. In *N. tabacum* and *A. comosus*, *CRC* was highly up-regulated in the nectaries (log2 ratio < 6). In a heatmap for each species, the transcript levels of different genes are shown (Supplementary Fig. S2). Only genes related to carbon metabolism, sugar transport or nectar secretion have been included here (SPS, INV, SUS, SUT, SWEET, STP, PIP, SNARE) A large number of genes with different expression levels were detected in the samples, so the dendrogram of the heatmap shows a total separation of the leaf and nectary samples for *N. tabacum* and *A. comosus* (Supplementary Fig. S2A, B).

### Gene Ontology (GO) enrichment analysis

Since there were no database entries for the two genomes (*N. tabacum* and *A. comosus*) in which the genes were associated with the GO terms, a corresponding GO background data file had to be created to perform the GO enrichment method. For *N. tabacum*, 41 % of the genes in the transcriptome had corresponding GO terms and for *A. comosus*, 42 %. Based on these annotations, GO enrichment was performed to verify the similarities or differences of each gene set with respect to its molecular function, cellular component or biological process. Two gene sets were used for each species. In one gene set, genes were up-regulated in the nectaries (log2 ratio > 1) and in the other gene set they were down-regulated in the nectaries, indicating that they were up-regulated in the leaves (log2 ratio < -1). As shown in Fig. 2, the enriched GO terms included molecular function, cellular component, and biological processes. In the GO enrichment analysis of *N. tabacum* nectaries, the “transporter activity” was the most enriched in the category of molecular functions, “endoplasmic reticulum” and “membrane” in the category of cellular components and “response to chemical”, “catabolic processes” and “flower development” in the category of biological processes (Fig. 2A). For the nectaries of *A. comosus*, the GO terms enriched in the three categories differed from those for *N. tabacum* (Fig. 2C). In *A. comosus*, “structural molecule activity” was most enriched in the category of molecular functions, “external encapsulating structure”, “cell wall” and “plasma membrane” were enriched in the category of cellular components and “anatomical structure development”, “cell growth” and “cell differentiation” were enriched in the category of biological processes. In both plant species, the up-regulated genes in nectaries and leaves differed considerably. Numerous genes with GO term up-regulated in leaves of *N. tabacum* and *A. comosus* were related to cellular components such as “chloroplasts and thylakoids” or to “photosynthesis” as a biological process (Fig. 2B, D).

**Figure 2:**
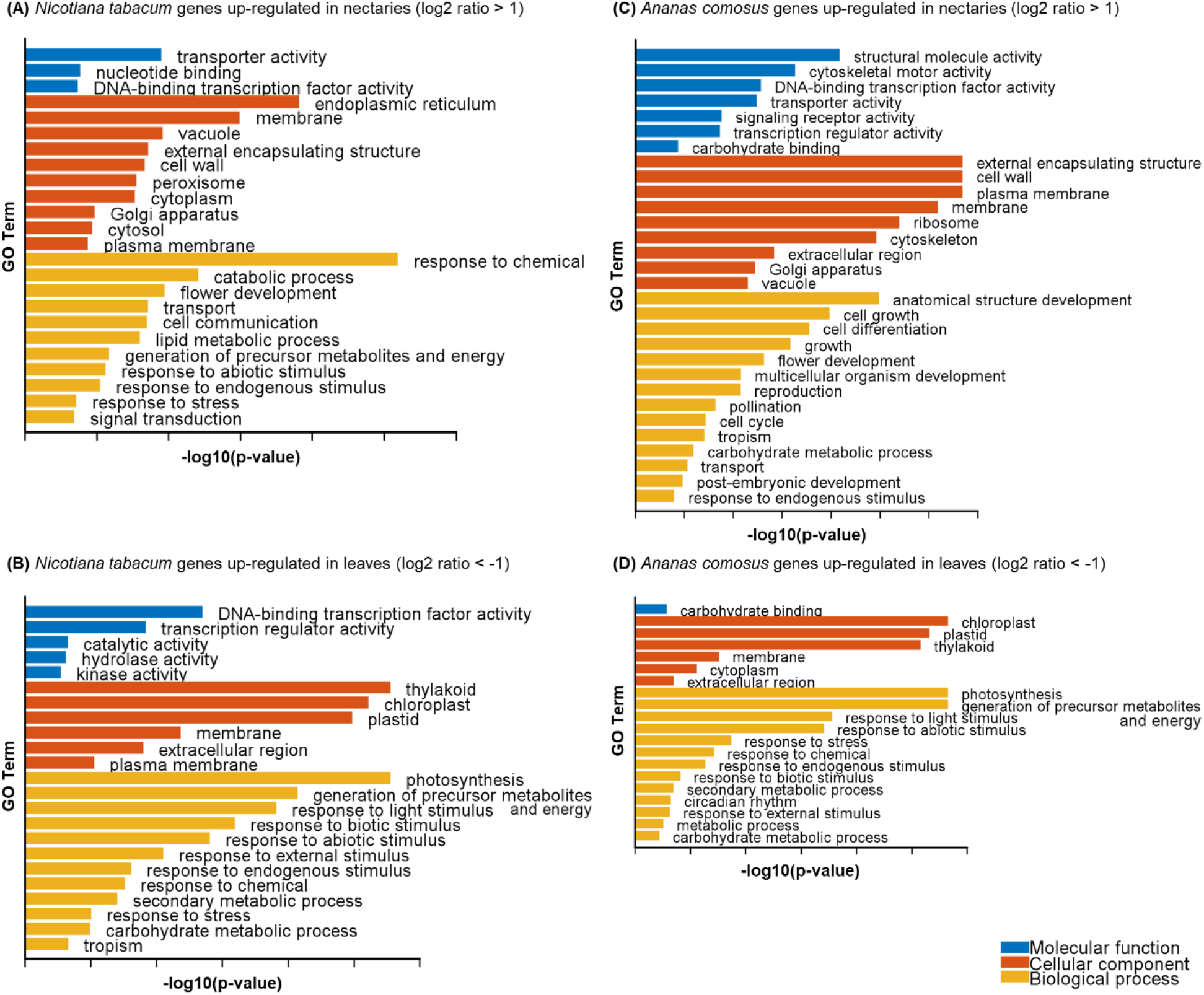
Gene Ontology (GO) enrichment analysis of *Nicotiana tabacum* and *Ananas comosus*. Gene ontology describes three aspects: molecular function (blue), cellular component (orange), and biological process (yellow). The genes or gene sets that were up-regulated according to these three aspects of the gene ontology. Different expressed genes (DEGs) in *Ananas comosus* (A, B) and *Nicotiana tabacum* (C, D) were used in the enrichment analyses, with a log2 ratio greater than 1 (A, C) and less than -1 (B, D). In the graphs the absolute confidences (-log10 adjusted *p* value) of the GO terms related to molecular function (blue), cellular component (orange), and biological process (yellow) were plotted.

### Expression levels of genes involved in sugar metabolism in nectaries and nectar secretion

After analyzing the entire transcriptomic dataset, various genes related to sugar metabolism in nectaries and nectar secretion were analyzed in more detail. The analyses included sucrose phosphate synthases, sucrose cleavage enzymes such as invertases and sucrose synthases; sugar transporters such as SWEETS; sucrose transporters (SUTs); hexose transporters (MST/HT) and aquaporins (plasma membrane intrinsic proteins, PIPs). Moreover, the expression of SNARE-genes was analyzed, because the corresponding proteins might be involved in exocytosis (granulocrine excretion). For each group of genes, the transcript expression in the nectaries relative to that in the leaves is shown as the log2 ratio (Supplementary Table S2, S3). If the log2 ratio is positive, the gene is up-regulated in the nectaries, and if it is negative, it is down-regulated in nectaries and thus up-regulated in leaves. For the comparison of leaves and nectaries, the transcription levels of the different genes are also shown in transcripts per million (TPM) for each species (Fig. 3-10). In general, TPM is normalized to the gene length and represents the abundance of a transcript within a population of sequenced transcripts. Moreover, phylogenetic analyses were performed for the selected genes (Supplementary Fig. S3-S10). For this purpose, in addition to the corresponding genes of *A. comosus* and *N. tabacum*, the corresponding genes of another dicot and another monocot plant, *Arabidopsis thaliana* and *Oryza sativa*, were included (Supplementary Table S4-S7).

**Figure 3:**
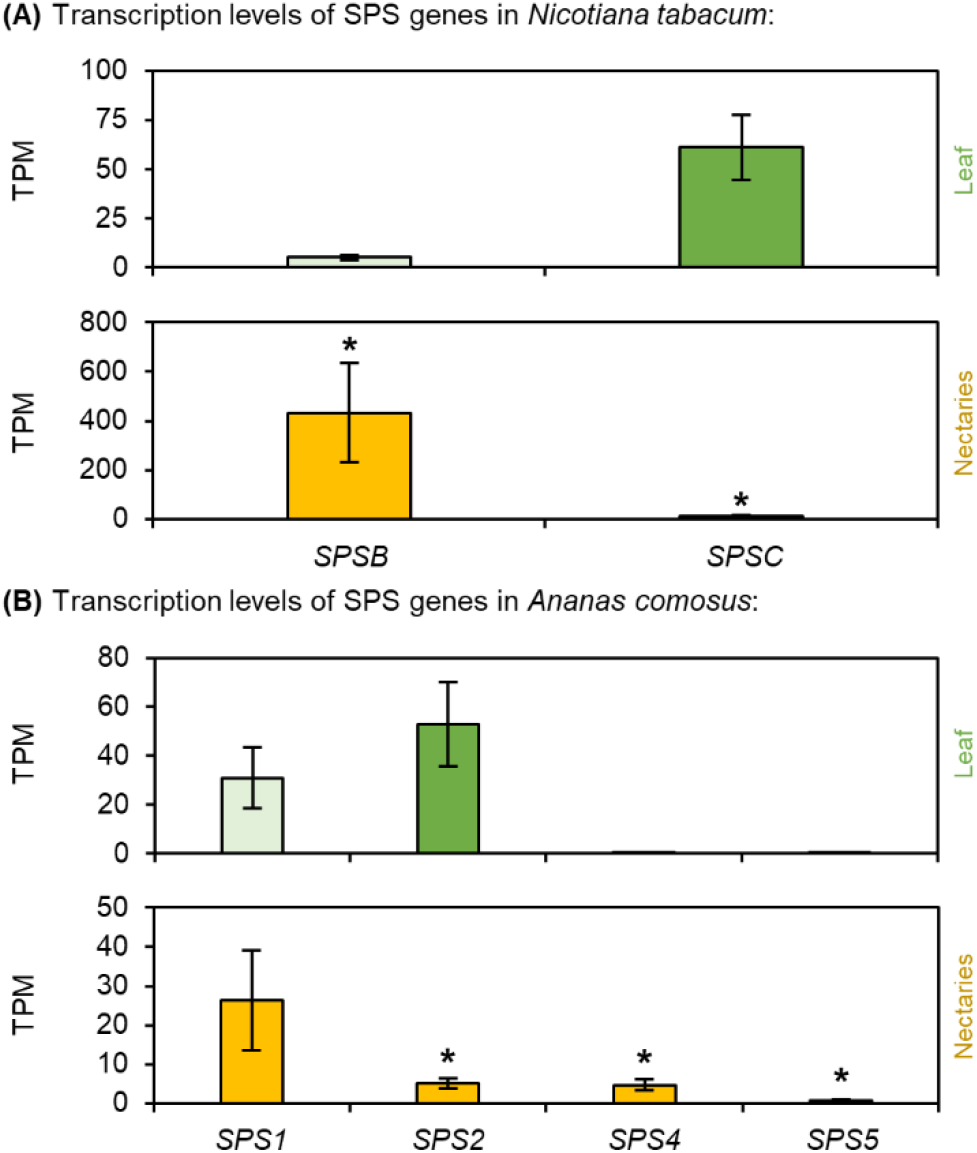
Transcription levels of sucrose-phosphate synthase (SPS) in *N. tabacum* and *A. comosus*. The results are separated for each species (A: *N. tabacum*; B: *A. comosus*). The two charts for each species (A, B) show the different TPM values for each SPS gene in the leaves and nectaries (mean **±** SD, n = 3). Error bars indicate the standard deviation. The significant difference between the genes in leaves and nectaries are indicated by asterisks (*t*-test, *p* < 0.05). These charts only allow the comparison of this gene group within the leaves or nectaries of a species.

### Sucrose-phosphate synthase (SPS)

Transcripts of two different SPS genes were found in *N. tabacum* (Fig. 3A). In the nectaries of *N. tabacum NtSPSB* had the highest TPM level and its expression was strongly up-regulated in the nectaries compared to the leaves (Fig. 3A). Transcripts of four SPS genes were found in *A. comosus* (Fig. 3 B), with *AcSPS1*, the closest ortholog to *AtSPS1F* (Supplementary Fig. S3), showing the highest TPM level. In *Arabidopsis, AtSPS1F* and *AtSPS2F* are essential for nectar production, because *sps1f/2f* mutants fail to secrete nectar [11]. However, the expression of *AcSPS1* was up-regulated in leaves compared to that in nectaries. In contrast, the expression of *AcSPS4* and *AcSPS5* was up-regulated in the nectaries, although both the TPM levels in the nectaries were lower than those in the *AcSPS1* (Fig. 3B; Supplementary Table S3).

### Invertase (INV)

Numerous invertase genes, including 10 neutral (NINV), three vacuolar (VINV), and three cell wall invertases (CWINV) were differentially expressed in *N. tabacum* (Fig. 4A). However, the TPM levels of several invertase genes were rather low, especially for CWINV (Fig. 4A). This also applies to *NtCWINV2*, the closest ortholog to *AtCWINV4* which is required for nectar production in *Arabidopsis* (Supplementary Fig. S4; Ruhlman et al., 2010). Compared to those in *N. tabacum*, a smaller number of invertase genes were expressed in *A. comosus*, one NINV, one VINV, and two CWINVs (Fig. 4B). The highest TPM level was found for *AcCWINV3* (Fig. 4B), which belongs to the same clade as *AtCWINV4* of *Arabidopsis*, but in the monocot group (Supplementary Fig. S4). In addition, *AcCWINV3* was the only invertase, whose expression was up-regulated in the nectaries compared to the leaves (Fig. 4 B; Supplementary Table S3).

**Figure 4:**
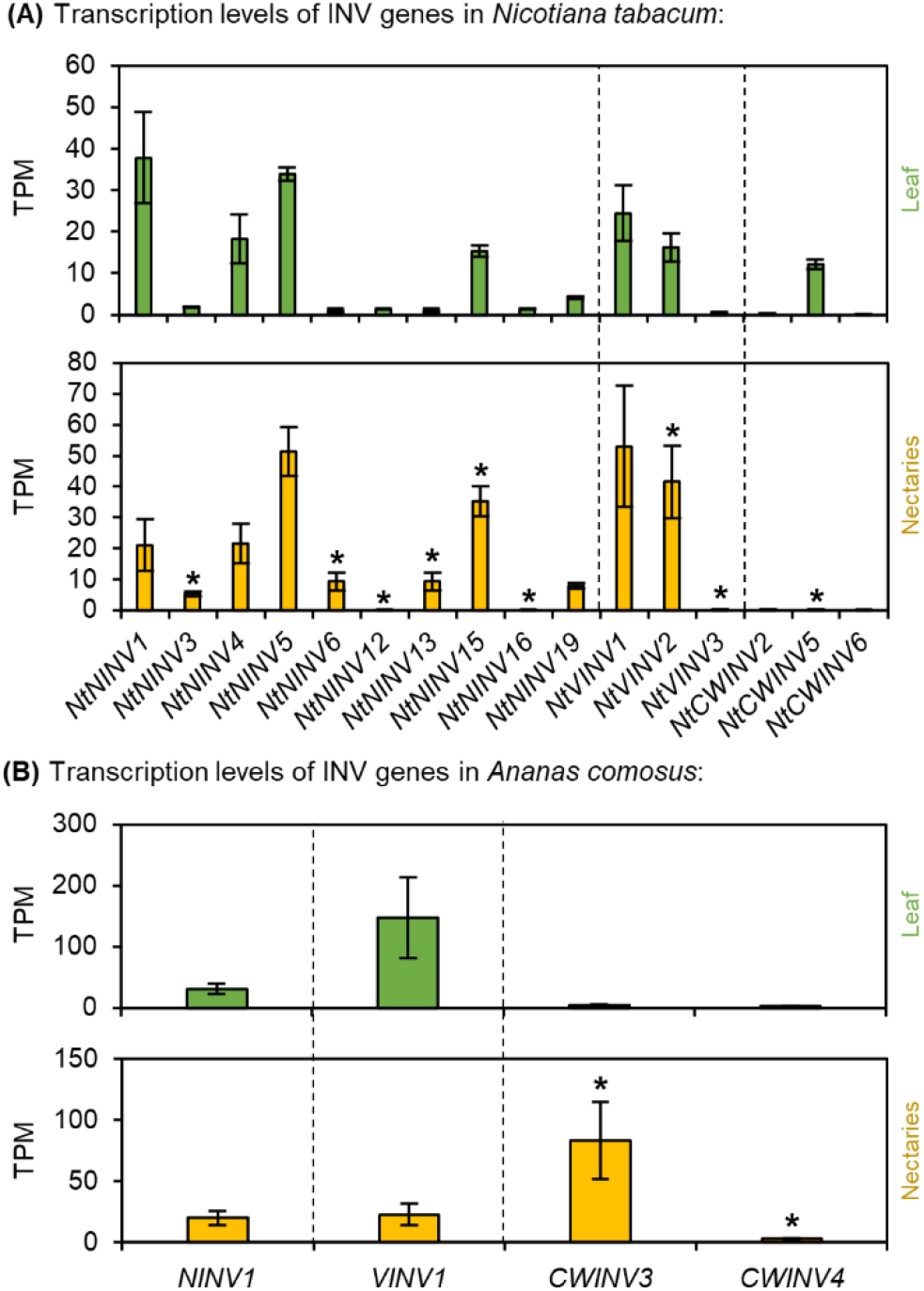
Transcription levels of different invertases (NINV, CWINV, VINV) in *N. tabacum* and *A. comosus*. The results are separated for each species (A: *N. tabacum*; B: *A. comosus*). The two charts for each species (A, B) show the different TPM values for each INV gene in the leaves and nectaries (mean **±** SD, n = 3). Error bars indicate the standard deviation. The significant difference between the genes in leaves and nectaries are indicated by asterisks (*t*-test, *p* < 0.05). These charts only allow the comparison of this gene group within the leaves or nectaries of a species.

### Sucrose synthase (SUS)

Phylogenetic analyses revealed that plant *SUS* genes can be divided into three separate clades, in which both dicots and monocots are represented (Supplementary Fig. S5). In *N. tabacum*, transcript abundance analysis showed that of the six *SUS* genes, almost all were up-regulated in the nectaries, in particular *NtSus1* was much more highly expressed in the nectaries than in the leaves (Fig. 5A; Supplementary Table S2). Furthermore, the TPM level of *NtSUS1* in the nectaries was at least 100-fold higher than the TPM levels of the other sucrose synthases (Fig. 5A). Transcripts of four *SUS* genes were found in *A. comosus* (Fig. 5B) and the highest TPM level was detected in the nectaries for *AcSUS1* followed by *AcSUS3* (Fig. 5B). However, both genes were up-regulated in leaves compared to nectaries but *AtSUS1.1*, which belongs to the same clade as *NtSUS1*, was up-regulated in nectaries (Fig. 5B; Supplementary Fig. S5; Supplementary Table S3).

**Figure 5:**
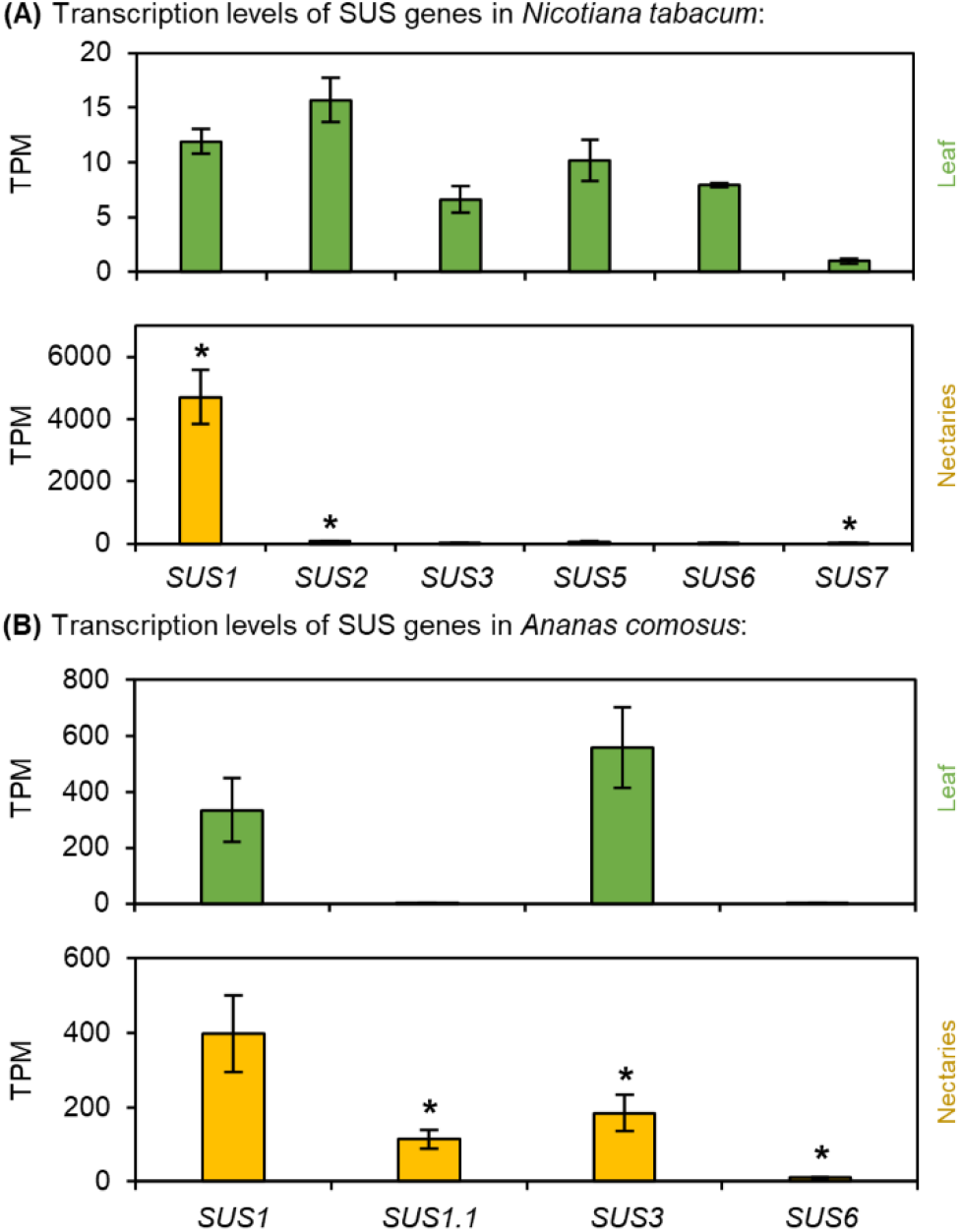
Transcription levels of sucrose synthase (SUS) in *N. tabacum* and *A. comosus*. The results are separated for each species (A: *N. tabacum*; B: *A. comosus*). The two charts for each species (A, B) show the different TPM values for each SUS gene in the leaves and nectaries (mean **±** SD, n = 3). Error bars indicate the standard deviation. The significant difference between the genes in leaves and nectaries are indicated by asterisks (*t*-test, *p* < 0.05). These charts only allow the comparison of this gene group within the leaves or nectaries of a species.

### SWEETs

Phylogenetic analyses of the SWEETs revealed that *A. thaliana* and *O. sativa*, like *N. tabacum* and *A. comosus* contain numerous SWEETs in different clades (Supplementary Fig. S6). Several SWEET genes were expressed in the leaves and nectaries of *N. tabacum* (Fig. 6A). *NtSWEET9* (clade III), the closest ortholog to *AtSWEET9* in *Arabidopsis,* which is essential for sucrose secretion in the nectaries of *A. thaliana* [11], showed the highest TPM value in nectaries of *N. tabacum* (Fig. 6A). Furthermore, the expression of *NtSWEET9* was strongly up-regulated in nectaries compared to leaves (Fig. 6A; Supplementary Table S2). *NtSWEET7* had the second highest TPM in the nectaries (clade II; Fig. 6A). *NtSWEET7* clustered with *AtSWEET7* from *Arabidopsis* and *CsSWEET7a* from *Cucumis sativus* (Supplementary Fig. S6).

**Figure 6:**
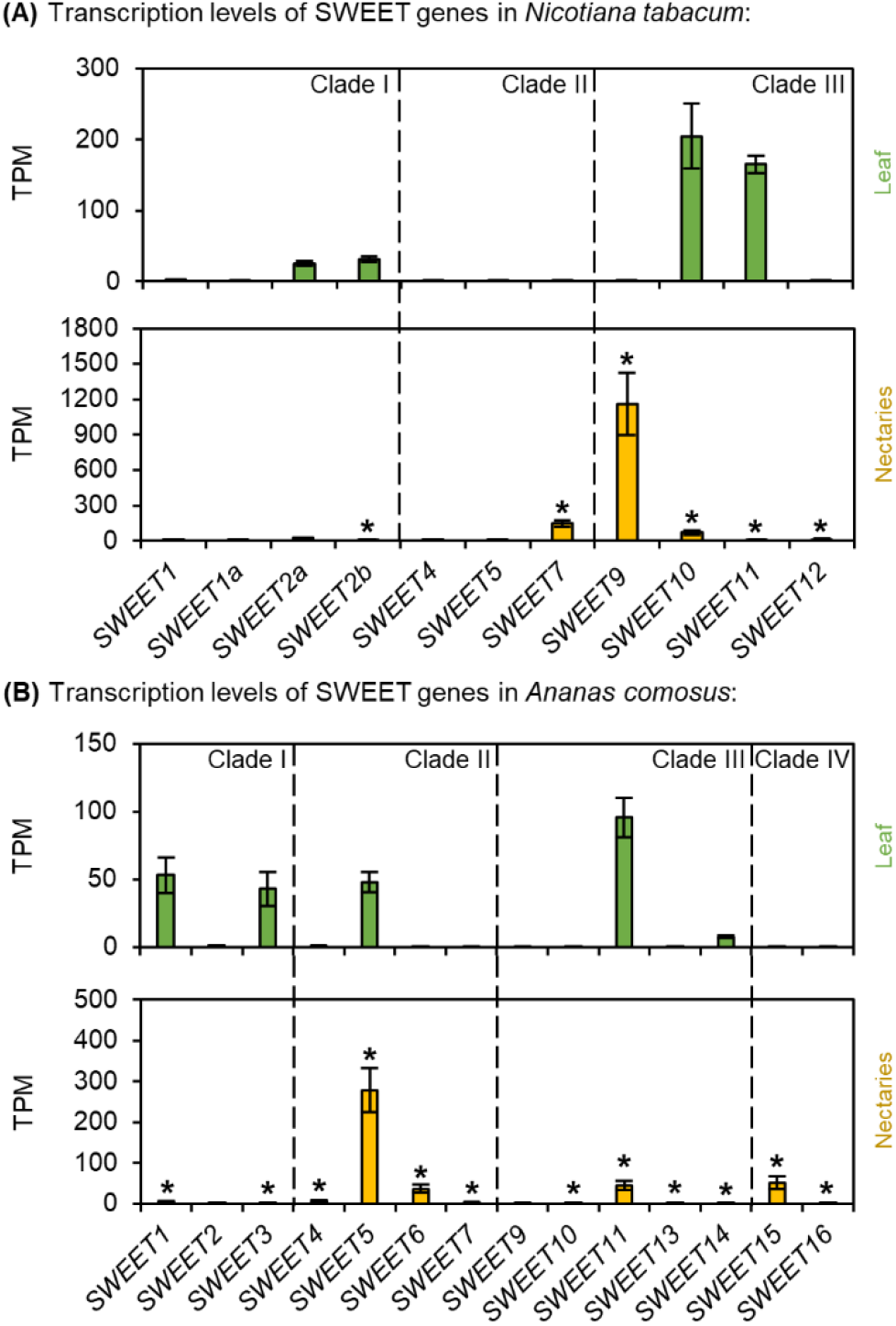
Transcription levels of sugars will eventually be exported transporters (SWEET) in *N. tabacum* and *A. comosus*. The results are separated for each species (A: *N. tabacum*; B: *A. comosus*). The two charts for each species (A, B) show the different TPM values for each SWEET gene in the leaves and nectaries (mean **±** SD, n = 3). Error bars indicate the standard deviation. The significant difference between the genes in leaves and nectaries are indicated by asterisks (*t*-test, *p* < 0.05). These charts only allow the comparison of this gene group within the leaves or nectaries of a species.

In *A. comosus*, numerous SWEET genes were also expressed, with about half of which were up-regulated in the nectaries, while the other half were up-regulated in the leaves (Fig. 6B; Supplementary Table S3). However, no orthologs of *AtSWEET9* or *NtSWEET9* (clade III) was found in *A. comosus* (Supplementary Fig. S6). Among the SWEETs of clade III, the highest TPM value was found for *AcSWEET11*, but this gene was up-regulated in the leaves but not in the nectaries (Fig. 6B; Supplementary Table S3). In the case of the SWEETs belonging to clade II, the expression of all SWEETs was up-regulated in the nectaries compared to the leaves (Fig. 6B; Supplementary Table S3). The TPM level of *AcSWEET5* (clade II) was the highest among the whole group of SWEETS in *A. comosus* (Fig. 6B). These results suggest that at least SWEET-mediated sucrose export (clade III SWEETs) may not be as important in the nectaries of *A. comosus* as in dicotyledons. Therefore, the expression levels of other sugar transporters, sucrose uptake transporters (SUTs) and monosaccharide transporters (STPs/HTs) were also tested.

### Sucrose transporters (SUTs)

Different types of sucrose uptake transporters (SUTs) were expressed in *N. tabacum* and *A. comosus* (Fig. 7; Supplementary Fig. S7). In *N. tabacum*, the highest TPM value was found for *NtSUT1* (type I), but the expression of *NtSUT1* was up-regulated in the leaves compared to the nectaries (Fig. 7A; Supplementary Table S2). In leaves of several dicotyledons, type I SUTs facilitate the active uptake of sucrose into the phloem [13]. In contrast, the expression of *NtSUT4* (type III) was up-regulated in nectaries of *N. tabacum* (Fig. 7A). Type III SUTs are functionally diverse and facilitate, for example, the active transport of sucrose from the vacuole into the cytoplasm [13].

**Figure 7:**
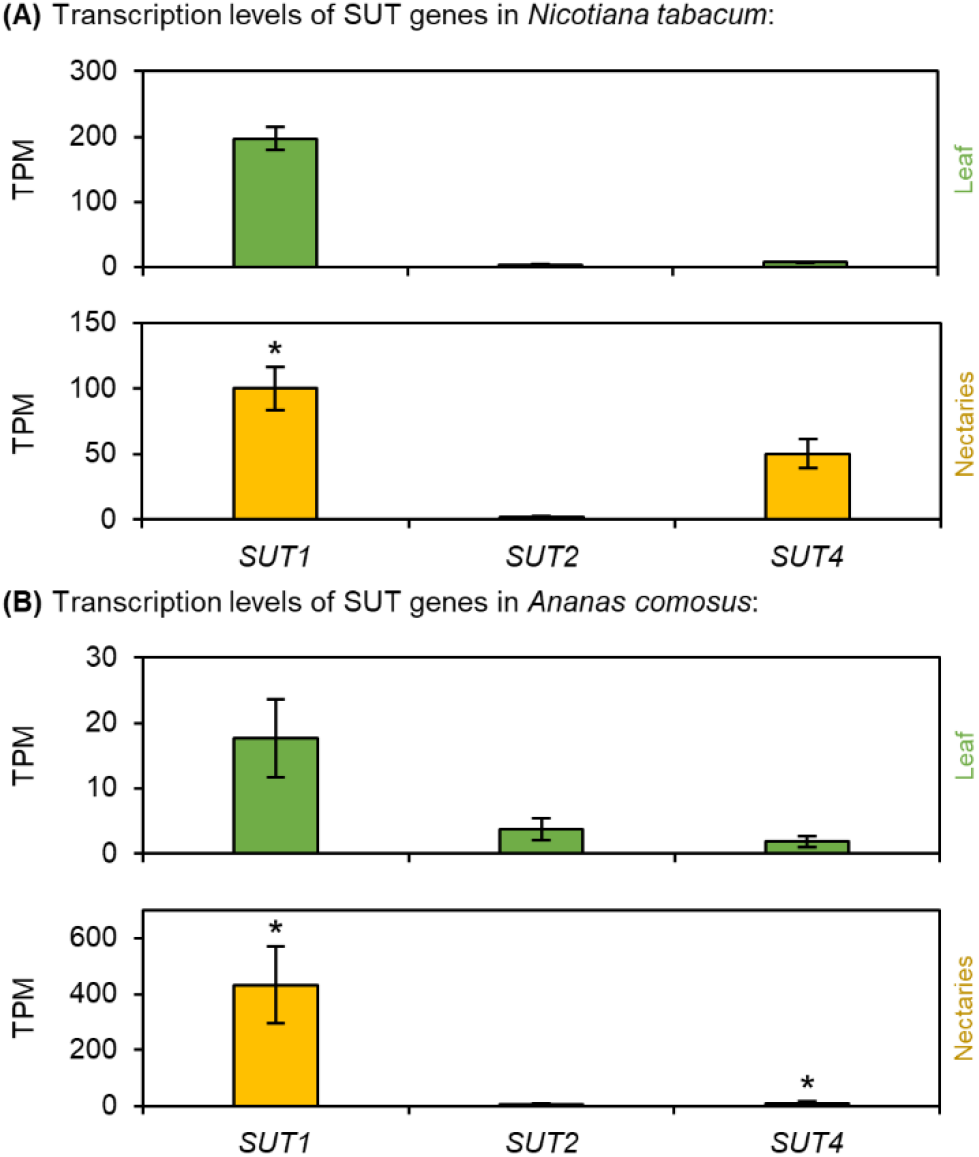
Transcription levels of sucrose uptake transporters (SUT) in *N. tabacum* and *A. comosus.* The results are separated for each species (A: *N. tabacum*; B: *A. comosus*). The two charts for each species (A, B) show the different TPM values for each SUT gene in the leaves and nectaries (mean **±** SD, n = 3). Error bars indicate the standard deviation. The significant difference between the genes in leaves and nectaries are indicated by asterisks (*t*-test, *p* < 0.05). These charts only allow the comparison of this gene group within the leaves or nectaries of a species.

In *A. comosus*, the highest TPM value was found for *AcSUT1* (type IIB, Supplementary Fig. S7) and the expression of *AcSUT1* was strongly up-regulated in the nectaries compared to the leaves (Fig. 7B; Supplementary Table S2). The monocot-specific type IIB contains SUTs that have a similar function to type I SUTs in dicots [13].

### Sugar transporter proteins (STPs)

In plants, a large number of STPs, which are H^+^/monosaccharide transporters that mediate the transport of hexoses across the plasma membrane in various plant cells, are expressed (Supplementary Fig. S8) [12]. In the nectaries of *N. tabacum*, the highest TPM values were found for *NtSTP23* and *NtSTP24* and the expression of both genes was up-regulated in the nectaries compared to the leaves (Fig. 7A; Supplementary Table S2). In *A. comosus*, the expression of more than half of the STPs was up-regulated in the nectaries compared to the leaves (Fig. 7B; Supplementary Table S3). However, the highest TPM value in the nectaries of *A. comosus* was found for *AcSTP1* (Fig. 7B).

### PIPs (plasma membrane intrinsic proteins)

Phylogenetic analyses revealed numerous PIP genes are represented in monocots and dicots, including *N. tabacum* and *A. comosus* (Supplementary Fig. S9). Moreover, in both *N. tabacum* and *A. comosus*, the expression of almost all PIPs was up-regulated in the nectaries compared to leaves (Fig. 9A, B; Supplementary Table S2, S3). The highest TPM value in the nectaries of *N. tabacum* was found for *NtPIP2-1*, followed by the closest homologs *NtPIP1-3* and *NtPIP2-3* (Fig. 9B; Supplementary Fig. S9). In the case of *A. comosus,* the highest transcript expression level was found for *AcPIP2-4*, followed by *AcPIP1-2* (Fig. 9B).

### *N*-ethylmaleimide-sensitive factor adaptor protein receptor (SNARE)-domain-containing proteins

In plants, SNAREs can be divided into five clades (Qa, Qb, Qc, Qb + Qc, and R) and their primary function is to mediate the fusion of vesicles with the target membrane, which is part of exocytosis. Phylogenetic analyses revealed that both dicots and monocots contain a larger number of SNARE genes (Supplementary Fig. S10). A large number of SNARE genes are expressed in both in *N. tabacum* and in *A. comosus*, and in both plant species the expression of up to three quarters of the SNARE genes was up-regulated in the nectaries (Fig. 10A, B; Supplementary Table S2, S3). In the case of *A. comosus* this is especially true for for *AcSYP123* (Fig. 10B; Supplementary Table S3). However, one difference between the two plant species was that transcripts of members of the clade Qb + Qc were only found in *A. comosus* (Fig. 10A, B; Supplementary Table S2, S3).

## Discussion

In *Arabidopsis*, which produces hexose-dominant nectar, several steps important for nectar secretion have been described, including sucrose synthesis by *AtSPS1F/2F* after the degradation of starch stored in nectaries, the export of sucrose by *AtSWEET9*, and sucrose hydrolysis during secretion by AtCWINV4 [11, 29, 58]. The nectar composition and the type of floral nectaries vary depending on the plant species. For this reason, different plant species will also use different modes of nectar production and secretion. To date, knowledge about nectar production in dicot species, especially rosids (e.g. *Arabidopsis, Brassica, Cucurbita*), is far greater than in monocot species. Therefore, in this study, the nectaries and nectar of *N. tabacum* (asterids) and *A. comosus* (monocot) were analyzed to test whether the mechanism described for *Arabidopsis* is also present in these plant species.

### Gene expression profiles and GO enrichment analysis of *N. tabacum* and *A. comosus*

Transcriptome sequencing (RNA-Seq) is an effective means of studying the molecular mechanisms of non-model plants and other organisms without whole-genome information [58–60]. In *N. tabacum* and *A. comosus*, the differences in gene expression between the nectaries and leaves were evaluated. Based on initial studies with volcano plots, principal component analysis (PCA), and expression heatmaps the data showed that the genes differed significantly between the nectaries and leaves (Supplementary Fig. S1, S2). More genes were up-regulated than down-regulated in the nectaries compared to the leaves. Due to their up-regulation in the nectaries, it can be assumed that a large number of genes are involved in flower growth and in the production and secretion of nectar, although this can vary depending on the flowering/nectar production stage [61, 62]. Furthermore, the strongly up-regulated expression of CRABS CLAW (*CRC*) in the nectaries demonstrated the veracity of the RNA-Seq data from the studied samples. The differential expression of the genes was also reflected in the GO enrichment analysis. Biological processes for flower development, reproduction, and pollination, for example, can be found in the nectaries using the GO terms, and photosynthesis, as well as response reactions to light stimulus, can be found in leaves (Supplementary Fig. 2). GO enrichment analyses in other species revealed similar results for the same tissue; for example, photosynthesis-related GO terms were found in the leaves of *Angelica glauca* [63]. Furthermore, expressed genes of a floral transcriptome of *Arabidopsis* could also be linked to flower development, reproduction and pollination [64].

### Nectar production and secretion in *N. tabacum*

#### Phloem unloading and sugar transport into the nectaries

A model about nectar production and secretion in *N. tabacum* is shown in Fig. 11A. During nectar secretion there is an increased need for carbohydrates in nectaries [9]. The phloem supplies the nectaries with sucrose either symplastically via plasmodesmata or sucrose is first transported into the apoplasm and then actively taken up into the nectaries. Apoplasmic unloading may be mediated by the reversal of SUTs, SWEET uniporters and/or other unidentified transporters [18, 65]. Unloaded sucrose can then either taken up into nectary cells by SUTs or hydrolyzed by CWINV to hexoses, which are taken up by hexose transporters (e.g. STPs). However, it is not yet clear which phloem unloading mode and sugar uptake mechanisms exist in nectaries.

In cucumber (*Cucumis sativus*), *CsSWEET7a* was highly expressed in flowers shortly before anthesis, and this protein was detected in the vascular tissues (phloem) of the receptacle and nectary [19]. These results, indicate that *CsSWEET7a* is involved in phloem unloading and sugar partitioning in the mentioned tissues [19]. In *N. tabacum,* the transcript expression levels of *NtSWEET7* and *NtSWEET4*, the closest orthologs to *CsSWEET7a* (Supplementary Fig. S6), were also much higher in the nectaries than in leaves (Fig. 6A; Supplementary Fig. S2). However, these SWEETs belong to clade II and transport mainly hexoses [66], whereas the phloem contains up to 1,000 millimolar sucrose and only low concentrations of hexoses [25]. In root tips, SUTs are located in the phloem, where they are expected to function in an efflux mode [65]. Sucrose transporters could also be involved in phloem unloading in nectaries [19], such as *NtSUT1*, whose transcripts were found in the nectaries (Fig. 7A; Supplementary Fig. S2). Some of the sucrose released to the apoplasm may be hydrolyzed into hexoses by CWINV [65]. The transport of hexoses from the apoplasm to the cytoplasm of nectary cells could be mediated by monosaccharide transporters (STPs). The transcript expression levels of several monosaccharide transporters (STPs), especially *NtSTP23/24*, were higher in nectaries than in leaves of *N. tabacum* (Fig. 8A).

**Figure 8:**
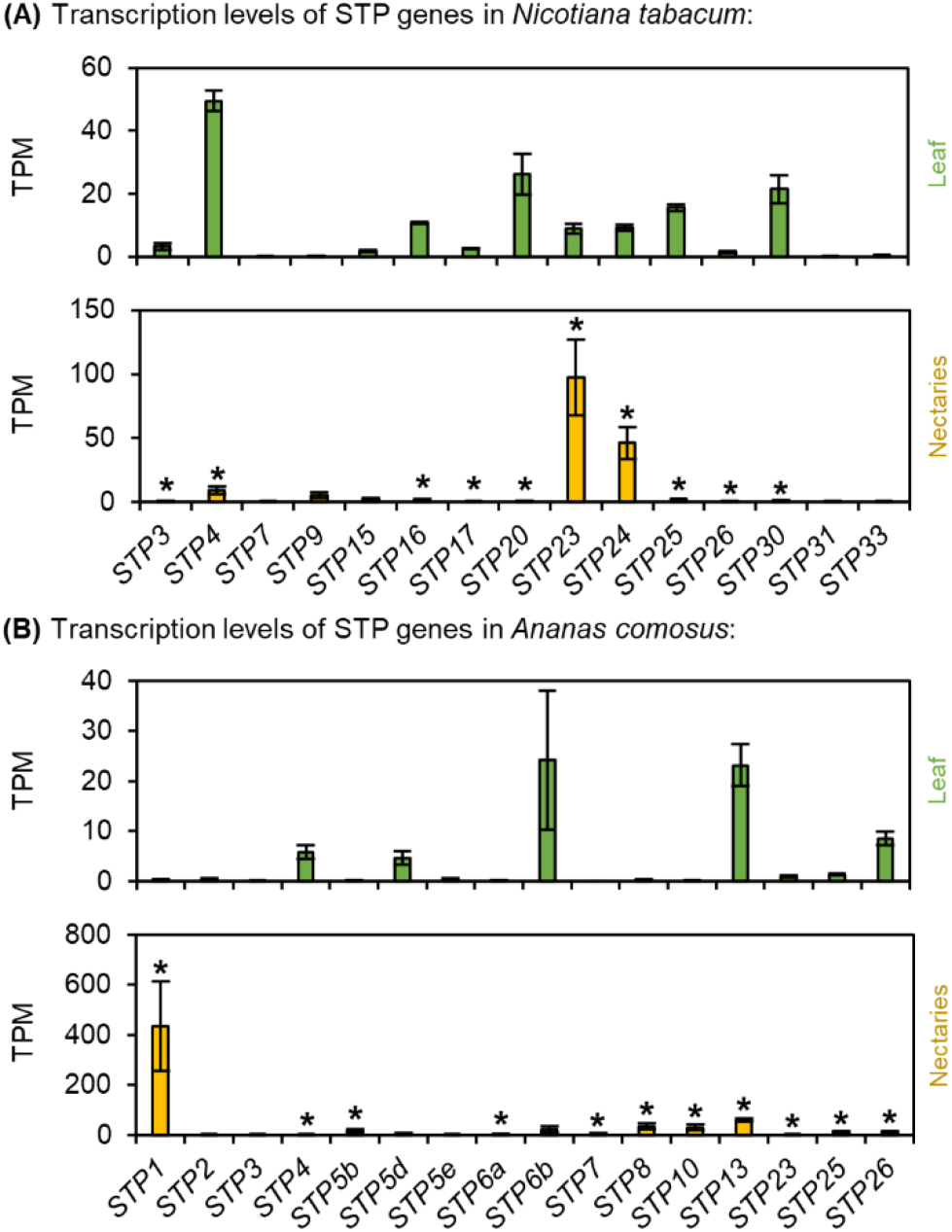
Transcription levels of sugar transport proteins (STP) in *N. tabacum* and *A. comosus*. The results are separated for each species (A: *N. tabacum*; B: *A. comosus*). The two charts for each species (A, B) show the different TPM values for each STP gene in the leaves and nectaries (mean **±** SD, n = 3). Error bars indicate the standard deviation. The significant difference between the genes in leaves and nectaries are indicated by asterisks (*t*-test, *p* < 0.05). These charts only allow the comparison of this gene group within the leaves or nectaries of a species.

### Nectary starch

Depending on the plant species, a certain portion of the sucrose delivered by the phloem is split into glucose and fructose in the nectaries [41, 42]. In some species, a portion of the glucose produced is temporarily stored in nectaries as starch, which is converted back to sugar during nectar secretion (Fig. 11A) [20, 23]. In *Arabidopsis*, sucrose-phosphate synthase genes (*SPS1F/2F*) are highly expressed in nectaries, and this enzyme may be involved in the re-synthesis of sucrose [58]. Moreover, sucrose synthesis was found to be important during nectar secretion, as *Arabidopsis* plants lacking sucrose phosphate synthases (*sps1f*/*2f* mutants) were unable to secrete nectar [11].

In *N. tabacum*, the starch content in the nectaries was much lower than in the leaves (Fig. 1E). A previous study revealed that the starch content in nectaries of day-flowering *Nicotiana* species such as *N. tabacum* was generally lower than that of night flowering species [41]. At night, the phloem transport of sucrose is reduced to about half of the daily rate [25, 67]. This means that less sucrose is probably supplied to the nectaries at night and that less sucrose is available directly for nectar production. Therefore, night flowering species may store more starch in the nectaries during the light period. At night, the stored starch is converted to sucrose, which is then used for nectar production. The reason may be that more sucrose is transported by the phloem and released to the nectaries during the day than at night [25, 67]. In day flowering species, such as *N. tabacum*, more phloem derived sucrose could be used directly for nectar sugar production. Nevertheless, starch degradation in nectaries also occurs in day-flowering species [22, 41]. The transcript expression level of *NtSPSB* was higher in the nectaries than in leaves of *N. tabacum* (Fig. 3B) and this enzyme might be involved in the re-synthesis of sucrose from starch in the nectaries.

### Sucrose cleavage enzymes in the nectaries

While the phloem sap of most plant species contains up to 1,000 millimolar sucrose and only small amounts of hexoses, the sucrose-to-hexoses ratio in the nectaries of *N. tabacum* was much lower (0.49; Table 1). This means, that a large portion of the sucrose has been split into glucose and fructose. Various cleavage enzymes can be involved in in this process, such as invertases or sucrose synthases (Fig. 11A). Several invertases were expressed in the nectaries of *N. tabacum*, but the expression level was not very high (Fig. 4A). Similar results were shown by another study [68], which analyzed the relative expression of cytosolic, vacuolar and cell wall invertases at different nectary developmental stages in *N. tabacum*. In contrast, the expression level of a sucrose synthase (*NtSUS1*) was very high in the nectaries of *N. tabacum* (Fig. 5A). There is evidence that sucrose synthases, rather than invertases, are the much more important cleavage enzymes in sink organs of several plant species [26] and SUS activity plays a crucial role in sink strength [45]. Pollen tubes of *N. tabacum* contain two isoforms of SUS and by immunohistochemical localization, one was found in the cytosol and associated with the plasma membrane, while the other was recognized only as being associated with the plasma membrane [69]. In *N. tabacum* nectaries, SUS may also be more involved in sucrose cleavage than invertases. In addition, the measured SUS activity in the nectaries of *N. tabacum* was higher than the activity of NINV (Table 2).

The sucrose-to-hexoses ratio in the nectar of *N. tabacum* was lower (0.26) than that in the nectaries (0.49), indicating that nectar contains more hexoses than sucrose compared to nectaries (Table 1). However, since the sugar composition in the phloem and in the nectaries differs much more than the composition in the nectaries and nectar, the sugar composition in the nectar is already largely determined by the metabolism in the nectaries and is only partly changed during secretion [41]. For several plant species with hexose-dominant nectar, such as *Arabidopsis thaliana* or *Brassica rapa*, it has been shown that cell wall invertases (CWINVs) are important for nectar secretion [29]. In addition, the hydrolysis of sucrose is also required to create an osmotic gradient large enough to allow water secretion [29]. The expression levels of various CWINVs in the nectaries of *N. tabacum* were rather low (Fig. 4A). In contrast, the CWINVs were still active at the same time (Table 2). In *N. tabacum* the expression of a cell wall invertase increased in the early flower developmental stages, but the expression decreased in the later stages [68]. Additionally, in the nectaries of *Cucurbita pepo*, the expression of *CpCWINV4* was high before anthesis but very low after anthesis [70]. During the different developmental stages of *N. tabacum* nectaries, the transcription of the CWINV genes is probably decreased before the enzyme activity declines. In addition to transcriptional regulation, the activity of the enzyme *in vivo* might also be regulated by several previously unknown mechanisms.

### Water secretion

A higher sugar concentration in nectar than in nectaries is necessary to create an osmotic gradient large enough to sustain water secretion [11]. It is assumed that plasma membrane intrinsic proteins (PIPs) may be involved in the rapid movement of water from nectary cells into nectar, but experimental evidence for this has yet to be provided [9]. However, in *Arabidopsis*, the expression of several aquaporins was increased compared to that in other tissues [58]. In *N. tabacum* the expression levels of several PIPs were also much higher in nectaries than in the leaves (Fig. 10A). Therefore, it can be assumed that these transporters may be involved in water secretion.

### SWEETs and sugar transport into nectar

In *N. tabacum* the sucrose concentration in nectar was only slightly higher than that in nectaries (Fig. 1C). Analyses of other day-flowering *Nicotiana* species also revealed that the sucrose concentration in the cytosol of nectary cells was slightly higher or similar to the sucrose concentration in the nectar [41]. At these sucrose concentrations, transport into the nectar can occur via facilitated diffusion transporters such as SWEET9 [11, 71]. The importance of SWEET9 for nectar secretion was demonstrated as *Arabidopsis* or *Nicotiana attenuata* mutants lacking *SWEET9* fail to produce nectar [11]. In *N. tabacum* nectaries, the transcript level of *NtSWEET9* was very high (Fig. 6A). Therefore, SWEET9 in *N. tabacum* is probably also involved in sucrose efflux from nectary cells into nectar. Since SWEET9 is a facilitated-diffusion transporter, it cannot secrete sugar at higher levels than those present in nectary cells [30]. Therefore, a sucrose concentration gradient must exist between nectary cells (symplasm) and nectar (apoplasm), and a high sucrose concentration must be maintained in the nectaries during nectar secretion [11]. Sucrose can either be synthesized in nectaries and/or delivered by the phloem. In addition, sugars can be released from the vacuole to increase the sugar concentration in the cytosol of nectary cells via sucrose transporters, e.g. *NtSUT4* (Fig. 11A). The transcript expression level of *NtSUT4* was high in the nectaries of *N. tabacum* (Fig. 7A).

### Sugar production and sugar secretion in the nectaries of *A. comosus*

The comparison of *A. comosus* and *N. tabacum* revealed numerous differences, both in the metabolite concentrations in nectar and nectaries as well as in the expression levels of various genes. A model about nectar production and secretion in *A. comosus* is shown in Fig. 11B.

### Nectary starch

Compared to *N. tabacum* the starch content in the nectaries of *A. comosus* was rather low (Fig. 1F). Microscopic analyses of *Ananas ananassoides* also showed that the starch reserves in the nectaries were almost completely hydrolyzed a few hours after anthesis [32]. Similar to various dicots [11], SPS might be involved in the re-synthesis of sucrose in *A. comosus*, as SPS was expressed in their nectaries, although not very strongly (Fig. 3B).

### Sucrose cleavage enzymes in the nectaries

Similar to other monocots, the sugar contained in the phloem sap of *A. comosus* probably consists almost exclusively of sucrose, which is delivered to the nectaries [17, 25]. Some of the sucrose must be cleaved there because the sucrose-to-hexoses ratio in nectaries was much lower than in phloem sap (Fig. 1D; Table 1) and the corresponding enzymes, invertases or sucrose synthases, were active in the nectaries. (Table 2). It should be noted that the *in vitro* activity of sucrose synthase was measured in the direction of sucrose degradation, but the enzyme could also be involved in sucrose synthesis under *in vivo* conditions in nectaries [72].

### SWEET9 is not present in the nectaries of monocots

The concentrations of both, hexoses and sucrose, in nectar were much higher than those in nectaries of *A. comosus* (Fig. 1D), which is consistent with the results for several other bromeliads [40]. However, it must also be noted that the measured metabolite concentration in the whole nectaries may differ from the concentrations in subdomains of the nectaries that are directly involved in nectar secretion [11]. In the nectaries of *A. comosus*, similar to those of *A. ananassoides*, a distinction can probably be made between the epithelium and nectary parenchyma [32], but the sugar concentrations in these cell types are not yet known.

While SWEET9 was found in members of the asterids (e.g. *Nicotiana*) and rosids (e.g. *Arabidopsis*), orthologs of SWEET9 appear to be absent in monocots such as *Musa acuminate* (banana) or *A. comosus*, both of which have septal nectaries (Fig. 6B) [11]. In contrast, in *A. comosus*, *SWEET11* (clade III), and *SWEET5*, *SWEET6* (both clade II), and *SWEET15* (clade IV) were highly expressed in the nectaries (Fig. 6B), suggesting that monocots may use other SWEETs for sugar efflux. The sequence of *AcSWEET11* was more similar to *AtSWEET9* than that of the other SWEETs and both clustered in clade III, whose members preferably mediate sucrose transport (Supplementary Fig. S6) [73]. However, the expression of *AcSWEET11* was up-regulated in the leaves and not in the nectaries, which speaks against a particular importance in the transport of sucrose from the nectaries into the nectar (Fig. 6B; Supplementary Table S3). In addition to a function in leaves, AcSWEET11 may play a critical role in fruit ripening in *A. comosus* [74]. In contrast, the expression of SWEETs of clade II were up-regulated in nectaries, for example *AcSWEET5* and *AcSWEET6*, which showed the highest transcript levels (Fig. 6B). Clade II members transport hexoses [73], making it unlikely that AcSWEET5 and AcSWEET6 are involved in sucrose transport in nectaries of

*A. comosus*. Clade IV members, such as AcSWEET15 can also be excluded, because they are located in the vacuole membrane and transport fructose [73]. In summary, SWEETs are probably not directly involved in the transport of sucrose from nectary cells into nectar in *A. comosus*.

In addition, since SWEETs function as facilitated diffusion transporters and the sugar concentration in nectar is probably higher than in nectaries of *A. comosus* (Fig. 1D), sugar secretion cannot be mediated exclusively by this type of transporters [30]. Therefore, other mechanisms must be involved in nectar secretion, such as the active transport of sugars [30]. To date, the possible involvement of active sugar transporters, for example sugar transport proteins (STPs; H^+^/monosaccharide symporters) or sucrose transporters (SUTs), in sugar secretion in nectaries has been postulated but not yet investigated [41, 75]. High transcript levels of several STPs, particularly *AcSTP1*, were found in the nectaries of *A. comosus* (Fig. 8B). *AcSTP1* expression was significantly higher in nectaries than in leaves. In the case of SUTs, *AcSUT1* showed the highest transcription level and *AcSUT1* was also much more expressed in nectaries than in leaves (Fig. 7B). SUTs are H^+^/sucrose symporters and for them to function in an efflux mode, the outward-directed chemical potential difference of sucrose across the plasma membrane must exceed the inward-directed proton motive forces [76]. Since the pH in the cytoplasm of nectary cells is probably neutral and slightly acidic in nectar [77, 78], and the sucrose concentration in nectar is also higher than in nectaries (Fig. 1D), it is unlikely that SUTs are involved in the active export of sucrose into the nectar. In conclusion, although active sugar export appears necessary, it is not fully known which transporters are involved in the secretion of sucrose from nectary cells into nectar of *A. comosus* and further experiments are required to answer this question (Fig. 11B).

### Sucrose cleavage enzymes in the nectaries

The sucrose-to-hexoses ratio in the nectar of *A. comosus* (0.30) was slightly higher than that in the nectaries (0.22; Table 1). Similar results were shown for other bromeliads with sucrose-rich nectar [40]. Therefore, the activity of sucrose cleavage enzymes in *A. comosus* appears to play a less important role in nectar secretion than in plant species with hexose-rich nectar, such as *N. tabacum* (Fig. 1C, D) or with hexose-dominant nectar, such as *Arabidopsis* [29]. This corresponds to a lower activity of CWINV in nectaries of *A. comosus* than in *N. tabacum* (Table 2). In species with hexose-dominant nectar, sucrose cleavage during nectar secretion may also be involved in creating a sufficient osmotic gradient to maintain water secretion [29]. Since sucrose cleavage during nectar secretion is probably rather low in *A. comosus*, other mechanisms are required to create a sufficient osmotic gradient. Similar to *N. tabacum*, the transcription levels of several PIPs were also much higher in nectaries than in leaves of *A. comosus* (Fig. 9; Supplementary Table S2, S3). However, whether these transporters are involved in water secretion remains to be investigated.

**Figure 9:**
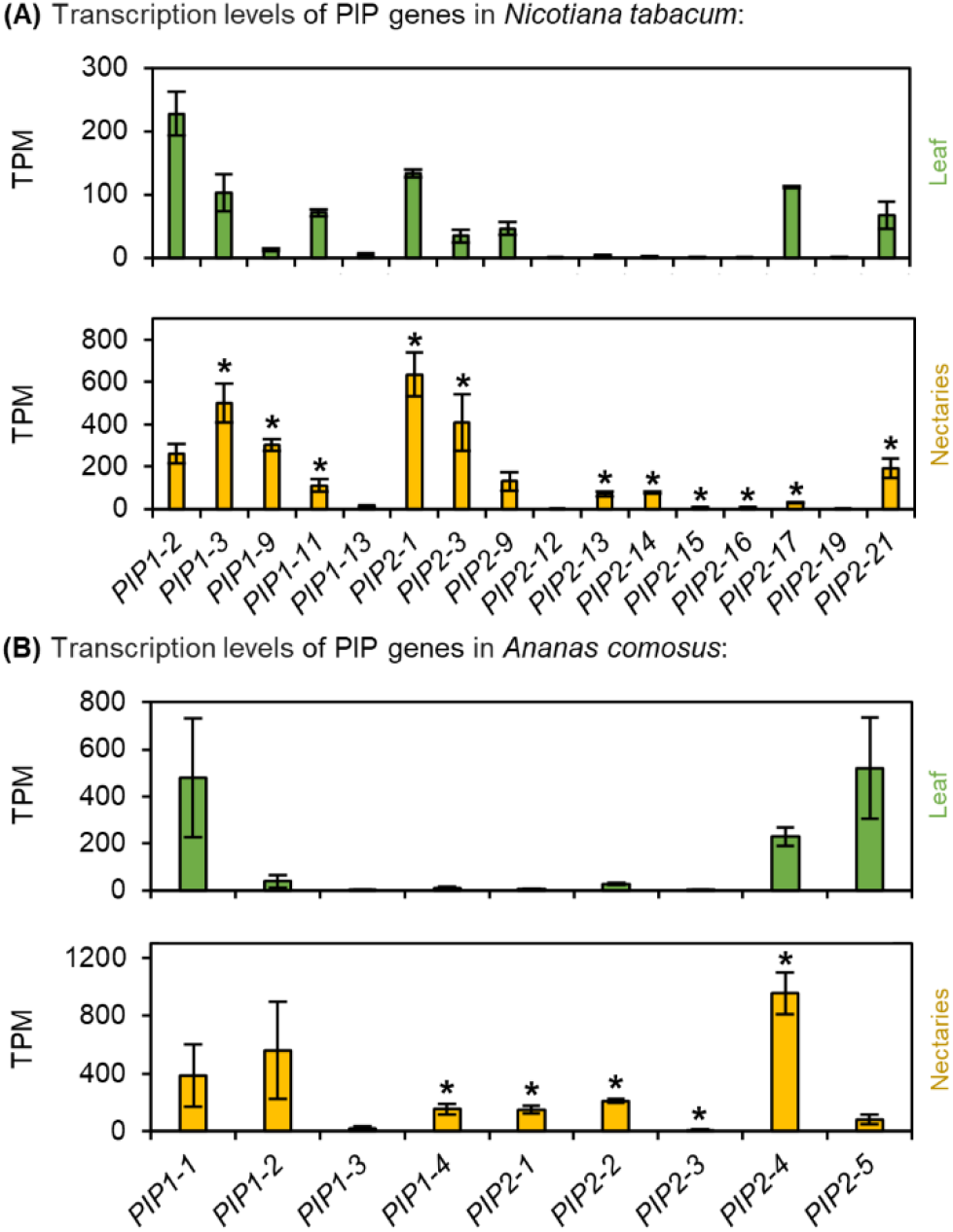
Transcription levels of aquaporins (PIP) in *N. tabacum* and *A. comosus*. The results are separated for each species (A: *N. tabacum*; B: *A. comosus*). The two charts for each species (A, B) show the different TPM values for each PIP gene in the leaves and nectaries (mean **±** SD, n = 3). Error bars indicate the standard deviation. The significant difference between the genes in leaves and nectaries are indicated by asterisks (*t*-test, *p* < 0.05). These charts only allow the comparison of this gene group within the leaves or nectaries of a species.

### Evidence of a granulocrine secretion mode in *A. comosus*

Based on ultrastructural analyses of the nectaries of *A. ananassoides,* it was hypothesized that metabolites are packaged into vesicles by the endoplasmic reticulum (ER) in the nectary cells [32]. The metabolites are released into the nectar after the vesicles fuse with the plasma membrane, the so-called granulocrine secretion [7]. The main function of SNAREs is to mediate the fusion of vesicles with the target membrane, which is part of exocytosis [33]. A large number of SNARE genes were expressed in *A. comosus*, and the expression of most SNARE genes was up-regulated in the nectaries, for example *AcSYP123* (Fig. 10B). Many genes whose GO term is related to the cell wall, such as “external encapsulating structure”, were also up-regulated in *A. comosus* nectaries (Fig. 2). Moreover, transcripts of members of the clade Qb + Qc were found only in *A. comosus* but not in *N. tabacum* (Fig. 10). However, further studies are needed to clarify whether SNAREs play a role in nectar secretion in *A. comosus.* Notably, the granulocrine type of secretion does not exclude involvement of plasma membrane transporters [10]. Moreover, the proposed models of nectar secretion are not necessarily mutually exclusive and it may be possible that different types of nectar secretion occur depending on the developmental stage or environmental conditions [79].

**Figure 10:**
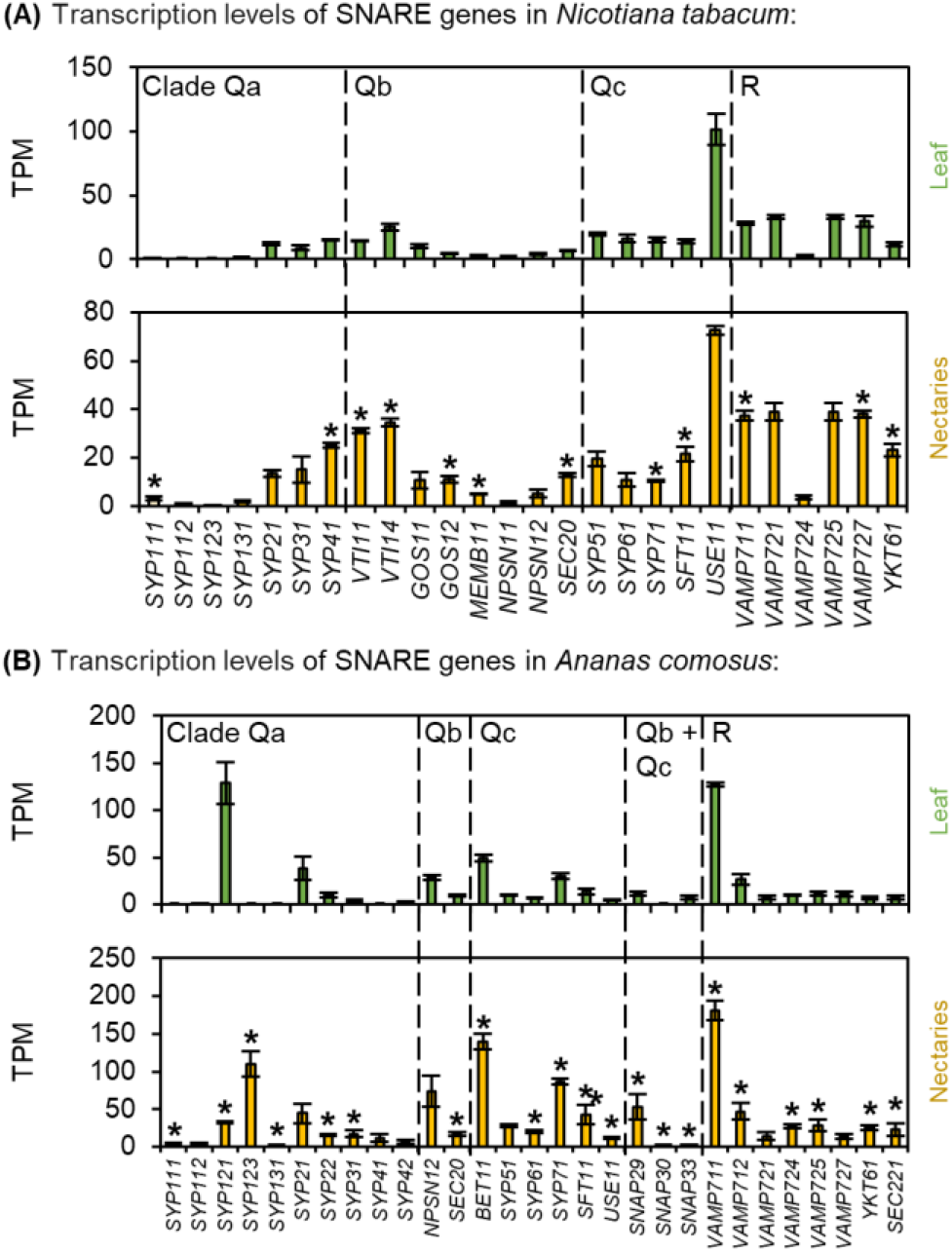
Transcription levels of soluble N-ethylmaleimide-sensitive-factor attachment receptor (SNARE)-domain-containing proteins in *N. tabacum* and *A. comosus*. The results are separated for each species (A: *N. tabacum*; B: *A. comosus*). The two charts for each species (A, B) show the different TPM values for each SNARE gene in the leaves and nectaries (mean **±** SD, n = 3). Error bars indicate the standard deviation. The significant difference between the genes in leaves and nectaries are indicated by asterisks (*t*-test, *p* < 0.05). These charts only allow the comparison of this gene group within the leaves or nectaries of a species.

**Figure 11:**
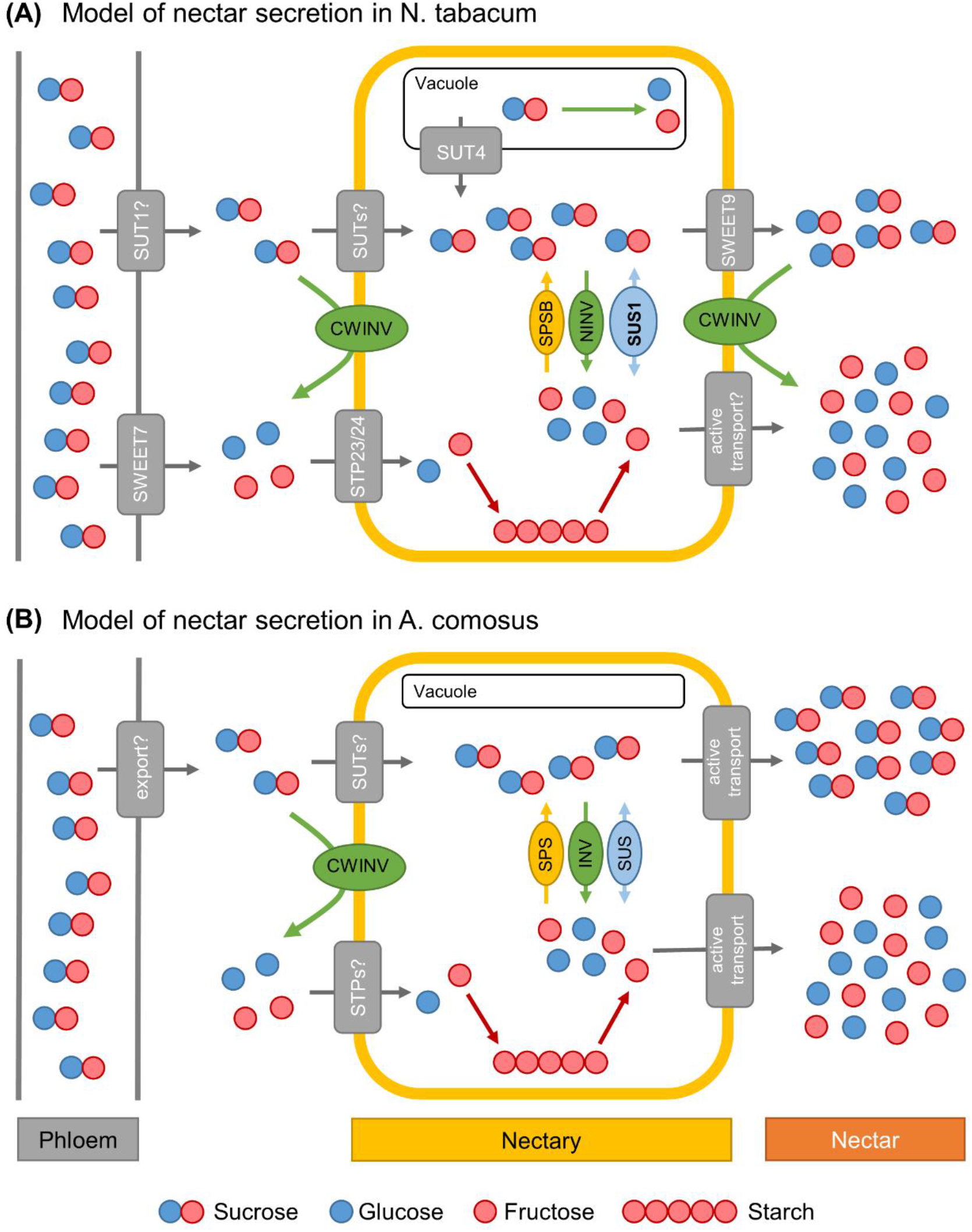
Models of nectar sugar secretion in *Nicotiana tabacum* (A) and *Ananas comosus* (B). SWEET: Sugar will eventually be exported transporter; SUT: Sucrose transporter; STP: Sugar transport protein; CWINV: Cell wall invertase; SPS: Sucrose-phosphate synthase; INV: Invertase; SUS: Sucrose synthase. The number of molecules corresponds to the concentration.

## Conclusion

The results show that the mechanisms of nectar production and secretion differ greatly between dicots and monocots. While many similarities were found between *N. tabacum* and *Arabidopsis*, major differences were found for *A. comosus*, including the absence of SWEET9 in monocots. Transcriptomic analyses of the nectaries of *A. comosus* enables many further investigations that will lead to a better understanding of nectar production in monocots. Furthermore, it may be the beginning of the investigation of the molecular mechanisms associated with granulocrine nectar secretion, e.g. the role of SNAREs in nectar secretion.

## Abbreviations

CAM: Crassulacean acid metabolism
CRC: CRABS CLAW
CWINV: Cell wall invertase
DEGs: Differentially expressed genes
ER: Endoplasmic reticulum
Fc: Fold change
GO: Gene Ontology
HPLC: High-performance liquid chromatography
INV: Invertase
NINV: Neutral/alkaline invertases
PC: Principal component
PCA: Principal component analysis
PIP: Plasma membrane intrinsic proteins
RNA-Seq: RNA Sequencing
SPS: Sucrose phosphate synthase
STP: Sugar transport protein
SUS: Sucrose synthase
SUT: Sucrose uptake transporter
SNARE: soluble N-ethylmaleimide-sensitive-factor attachment receptor (SNARE)-domain-containing protein
SWEET: Sugars will eventually be exported transporter
TPM: Transcript per million
t-SNARE: Target membrane-associated SNARE
UDP: Uridine-5-diphosphate
VINV: Vacuolar invertases
v-SNARE: vesicle-associated SNARE.

## Supporting Information

**Supplementary Figure S1**: Volcano plot, PCA plot, and Venn diagram of the transcriptomic data of *N. tabacum* and *A. comosus*.

**Supplementary Figure S2**: Expression heatmaps of the *Nicotiana tabacum* (A) and *Ananas comosus* (B) genes.

**Supplementary Figure S3**: Phylogenetic analysis of selected sucrose phosphate synthases (SPS).

**Supplementary Figure S4**: Phylogenetic analysis of selected cell wall invertases (CWINV).

**Supplementary Figure S5**: Phylogenetic analysis of selected sucrose synthases (SUS).

**Supplementary Figure S6**: Phylogenetic analysis of selected sugars will eventually be exported transporters (SWEET).

**Supplementary Figure S7**: Phylogenetic analysis of selected sucrose transporters (SUT).

**Supplementary Figure S8**: Phylogenetic analysis of selected sugar transport proteins (STP).

**Supplementary Figure S9**: Phylogenetic analysis of selected plasma membrane intrinsic proteins (PIP).

**Supplementary Figure S10:** Phylogenetic analysis of selected soluble N-ethylmaleimide-sensitive-factor attachment receptor (SNARE)-domain-containing proteins.

**Supplementary Table S1:** Water content in nectaries and leaves, expressed as a percentage of fresh weight.

**Supplementary Table S2:** Differential expression log2 ratios of different genes in *Nicotiana tabacum*.

**Supplementary Table S3:** Differential expression log2 ratios of different genes in *Ananas comosus*.

**Supplementary Table S4:** Gene IDs of *Arabidopsis thaliana* genes.

**Supplementary Table S5:** Gene IDs of *Nicotiana tabacum* genes.

**Supplementary Table S6:** Gene IDs of *Oryza sativa* genes.

**Supplementary Table S7:** Gene IDs of *Ananas comosus* genes.

## References

1. Brandenburg A, Dell’Olivo A, Bshary R, Kuhlemeier C. The sweetest thing: advances in nectar research. Curr Opin Plant Biol. 2009;12:486–90. doi:10.1016/j.pbi.2009.04.002.

2. González-Teuber M, Heil M. The role of extrafloral nectar amino acids for the preferences of facultative and obligate ant mutualists. J Chem Ecol. 2009;35:459–68. doi:10.1007/s10886-009-9618-4.

3. Nicolson SW. Sweet solutions: nectar chemistry and quality. Philos Trans R Soc Lond, B, Biol Sci. 2022;377:20210163. doi:10.1098/rstb.2021.0163.

4. Baker HG, Baker I. Floral nectar sugar constituents in relation to pollinator type. In: Jones CE, Little RJ, editors. Handbook of Experimental Pollination Biology. New York: Van Nostrand Reinhold; 1983b. p. 117–41. doi:10.2307/2443763.

5. Tiedge K, Lohaus G. Nectar sugars and amino acids in day- and night-flowering *Nicotiana* species are more strongly shaped by pollinators’ preferences than organic acids and inorganic ions. PLoS ONE. 2017;12:e0176865. doi:10.1371/journal.pone.0176865.

6. Göttlinger T, Schwerdtfeger M, Tiedge K, Lohaus G. What do nectarivorous bats like? Nectar composition in Bromeliaceae with special emphasis on bat-pollinated species. Front Plant Sci. 2019;10:205. doi:10.3389/fpls.2019.00205.

7. Fahn A. Ultrastructure of nectaries in relation to nectar secretion. Am. J. Bot. 1979b;66:977–85. doi:10.1002/j.1537-2197.1979.tb06309.x.

8. Pacini E, Nepi M. Nectar production and presentation. In: Nicolson SW, Nepi M, Pacini E, editors. Nectaries and nectar. Dordrecht: Springer Netherlands; 2007. p. 167–214. doi:10.1007/978-1-4020-5937-7_4.

9. Roy R, Schmitt AJ, Thomas JB, Carter CJ. Review: Nectar biology: From molecules to ecosystems. Plant Sci. 2017;262:148–64. doi:10.1016/j.plantsci.2017.04.012.

10. Kram BW, Carter CJ. *Arabidopsis thaliana* as a model for functional nectary analysis. Sex Plant Reprod. 2009;22:235–46. doi:10.1007/s00497-009-0112-5.

11. Lin IW, Sosso D, Chen L-Q, Gase K, Kim S-G, Kessler D, et al. Nectar secretion requires sucrose phosphate synthases and the sugar transporter SWEET9. Nature. 2014;508:546–9. doi:10.1038/nature13082.

12. Büttner M. The monosaccharide transporter(-like) gene family in *Arabidopsis*. FEBS Letters. 2007;581:2318–24. doi:10.1016/j.febslet.2007.03.016.

13. Sauer N. Molecular physiology of higher plant sucrose transporters. FEBS Letters. 2007;581:2309–17. doi:10.1016/j.febslet.2007.03.048.

14. Peng D, Gu X, Xue L-J, Leebens-Mack JH, Tsai C-J. Bayesian phylogeny of sucrose transporters: ancient origins, differential expansion and convergent evolution in monocots and dicots. Front Plant Sci. 2014;5:615. doi:10.3389/fpls.2014.00615.

15. Chen L-Q, Hou B-H, Lalonde S, Takanaga H, Hartung ML, Qu X-Q, et al. Sugar transporters for intercellular exchange and nutrition of pathogens. Nature. 2010;468:527–32. doi:10.1038/nature09606.

16. Chen L-Q, Qu X-Q, Hou B-H, Sosso D, Osorio S, Fernie AR, Frommer WB. Sucrose efflux mediated by SWEET proteins as a key step for phloem transport. Science. 2012;335:207–11. doi:10.1126/science.1213351.

17. Lohaus G, Schwerdtfeger M. Comparison of sugars, iridoid glycosides and amino acids in nectar and phloem sap of *Maurandya barclayana, Lophospermum erubescens*, and *Brassica napus*. PLoS ONE. 2014;9:e87689. doi:10.1371/journal.pone.0087689.

18. Borghi M, Fernie AR. Floral metabolism of sugars and amino acids: Implications for pollinators’ preferences and seed and fruit Set. Plant Physiol. 2017;175:1510–24. doi:10.1104/pp.17.01164.

19. Li Y, Liu H, Yao X, Sun L, Sui X. The role of sugar transporter CsSWEET7a in apoplasmic phloem unloading in receptacle and nectary during cucumber anthesis. Front Plant Sci. 2021;12:758526. doi:10.3389/fpls.2021.758526.

20. Ren G, Healy RA, Klyne AM, Horner HT, James MG, Thornburg RW. Transient starch metabolism in ornamental tobacco floral nectaries regulates nectar composition and release. Plant Sci. 2007;173:277–90. doi:10.1016/j.plantsci.2007.05.008.

21. Ning X, Tang T, Wu H. Relationship between the morphological structure of floral nectaries and the formation, transport, and secretion of nectar in lychee. Trees. 2017;31:1–14. doi:10.1007/s00468-016-1504-4.

22. Solhaug EM, Johnson E, Carter CJ. Carbohydrate metabolism and signaling in squash nectaries and nectar throughout floral maturation. Plant Physiol. 2019a;180:1930–46. doi:10.1104/pp.19.00470.

23. Nepi M, Ciampolini F, Pacini E. Development and ultrastructure of *Cucurbita pepo* nectaries of male flowers. Ann Bot. 1996;78:95–104.

24. Horner HT, Healy RA, Ren G, Fritz D, Klyne A, Seames C, Thornburg RW. Amyloplast to chromoplast conversion in developing ornamental tobacco floral nectaries provides sugar for nectar and antioxidants for protection. Am. J. Bot. 2007;94:12–24. doi:10.3732/ajb.94.1.12.

25. Lohaus G. Review primary and secondary metabolites in phloem sap collected with aphid stylectomy. J Plant Physiol. 2022;271:153645. doi:10.1016/j.jplph.2022.153645.

26. Stein O, Granot D. An overview of sucrose synthases in plants. Front Plant Sci. 2019;10:95. doi:10.3389/fpls.2019.00095.

27. Sturm A. Invertases. Primary structures, functions, and roles in plant development and sucrose partitioning. Plant Physiol. 1999;121:1–8. doi:10.1104/pp.121.1.1.

28. Roitsch T, González M-C. Function and regulation of plant invertases: Sweet sensations. Trends Plant Sci. 2004;9:606–13. doi:10.1016/j.tplants.2004.10.009.

29. Ruhlmann JM, Kram BW, Carter CJ. *CELL WALL INVERTASE 4* is required for nectar production in *Arabidopsis*. J Exp Bot. 2010;61:395–404. doi:10.1093/jxb/erp309.

30. Kim J-Y, Loo EP-I, Pang TY, Lercher M, Frommer WB, Wudick MM. Cellular export of sugars and amino acids: Role in feeding other cells and organisms. Plant Physiol. 2021;187:1893–914. doi:10.1093/plphys/kiab228.

31. Maurel C, Boursiac Y, Luu D-T, Santoni V, Shahzad Z, Verdoucq L. Aquaporins in Plants. Physiol Rev. 2015;95:1321–58. doi:10.1152/physrev.00008.2015.

32. Stahl JM, Nepi M, Galetto L, Guimarães E, Machado SR. Functional aspects of floral nectar secretion of Ananas ananassoides, an ornithophilous bromeliad from the Brazilian savanna. Ann Bot. 2012;109:1243–52. doi:10.1093/aob/mcs053.

33. Lipka V, Kwon C, Panstruga R. SNARE-ware: the role of SNARE-domain proteins in plant biology. Annu Rev Cell Dev Biol. 2007;23:147–74. doi:10.1146/annurev.cellbio.23.090506.123529.

34. Gu X, Brennan A, Wei W, Guo G, Lindsey K. Vesicle transport in plants: A revised phylogeny of SNARE proteins. Evol Bioinform Online. 2020;16:1176934320956575. doi:10.1177/1176934320956575.

35. Rahman AMA, Kumar VS. Estimation of the pineapple genome size by using quantitative real-time polymerase chain reaction. In: 9^th^ Malaysia Genetics Congress; 2011.

36. Sierro N, Battey JND, Ouadi S, Bakaher N, Bovet L, Willig A, et al. The tobacco genome sequence and its comparison with those of tomato and potato. Nat Commun. 2014;5:3833. doi:10.1038/ncomms4833.

37. Benzing DH. Bromeliaceae: Profile of an adaptive radiation. Cambridge: Cambridge University Press; 2000.

38. Bernardello G. A systematic survey of floral nectaries. In: Nicolson SW, Nepi M, Pacini E, editors. Nectaries and nectar. Dordrecht: Springer Netherlands; 2007. p. 19–128. doi:10.1007/978-1-4020-5937-7_2.

39. Sajo MG, Rudall PJ, Prychid CJ. Floral anatomy of Bromeliaceae, with particular reference to the evolution of epigyny and septal nectaries in commelinid monocots. Plant Syst. Evol. 2004. doi:10.1007/s00606-002-0143-0.

40. Göttlinger T, Lohaus G. Comparative analyses of the metabolite and ion concentrations in nectar, nectaries, and leaves of 36 bromeliads with different photosynthesis and pollinator types. Front Plant Sci. 2022;13:987145. doi:10.3389/fpls.2022.987145.

41. Tiedge K, Lohaus G. Nectar sugar modulation and cell wall invertases in the nectaries of day- and night-flowering *Nicotiana*. Front Plant Sci 2018. doi:10.3389/fpls.2018.00622.

42. Göttlinger T, Lohaus G. Influence of light, dark, temperature and drought on metabolite and ion composition in nectar and nectaries of an epiphytic bromeliad species (*Aechmea fasciata*). Plant Biol. 2020;22:781–93. doi:10.1111/plb.13150.

43. Lunn J, Hatch M. Primary partitioning and storage of photosynthate in sucrose and starch in leaves of C4 plants. Planta. 1995;197:385–91. doi:10.1007/BF00202661.

44. Heineke D, Sonnewald U, Bussis D, Gunter G, Leidreiter K, Wilke I, et al. Apoplastic expression of yeast-derived invertase in potato: Effects on photosynthesis, leaf solute composition, water relations, and tuber composition. Plant Physiol. 1992;100:301–8. doi:10.1104/pp.100.1.301.

45. Zrenner R, Salanoubat M, Willmitzer L, Sonnewald U. Evidence of the crucial role of sucrose synthase for sink strength using transgenic potato plants (*Solanum tuberosum* L.). Plant J. 1995;7:97–107. doi:10.1046/j.1365-313x.1995.07010097.x.

46. Morell M, Copeland L. Sucrose synthase of soybean nodules. Plant Physiol. 1985;78:149–54. doi:10.1104/pp.78.1.149.

47. Chang S, Puryear J, Cairney J. A simple and efficient method for isolating RNA from pine trees. Plant Mol Biol Rep. 1993;11:113–6. doi:10.1007/BF02670468.

48. Martin M. Cutadapt removes adapter sequences from high-throughput sequencing reads. EMBnet j. 2011;17:10. doi:10.14806/ej.17.1.200.

49. Langmead B, Salzberg SL. Fast gapped-read alignment with Bowtie 2. Nat Methods. 2012;9:357–9. doi:10.1038/nmeth.1923.

50. Ming R, VanBuren R, Wai CM, Tang H, Schatz MC, Bowers JE, et al. The pineapple genome and the evolution of CAM photosynthesis. Nat Genet. 2015;47:1435–42. doi:10.1038/ng.3435.

51. Edwards KD, Fernandez-Pozo N, Drake-Stowe K, Humphry M, Evans AD, Bombarely A, et al. A reference genome for *Nicotiana tabacum* enables map-based cloning of homeologous loci implicated in nitrogen utilization efficiency. BMC Genom. 2017;18:448. doi:10.1186/s12864-017-3791-6.

52. Love MI, Huber W, Anders S. Moderated estimation of fold change and dispersion for RNA-seq data with DESeq2. Genome Biol. 2014;15:550. doi:10.1186/s13059-014-0550-8.

53. Huerta-Cepas J, Szklarczyk D, Heller D, Hernández-Plaza A, Forslund SK, Cook H, et al. eggNOG 5.0: a hierarchical, functionally and phylogenetically annotated orthology resource based on 5090 organisms and 2502 viruses. Nucleic Acids Res. 2019;47:D309–D314. doi:10.1093/nar/gky1085.

54. Cantalapiedra CP, Hernández-Plaza A, Letunic I, Bork P, Huerta-Cepas J. eggNOG-mapper v2: Functional annotation, orthology assignments, and domain prediction at the metagenomic scale. Mol Biol Evol. 2021;38:5825–9. doi:10.1093/molbev/msab293.

55. Chen C, Wu Y, Li J, Wang X, Zeng Z, Xu J, et al. TBtools-II: A “one for all, all for one” bioinformatics platform for biological big-data mining. Mol Plant. 2023;16:1733–42. doi:10.1016/j.molp.2023.09.010.

56. Tamura K, Stecher G, Peterson D, Filipski A, Kumar S. MEGA6: Molecular evolutionary genetics analysis version 6.0. Mol Biol Evol. 2013;30:2725–9. doi:10.1093/molbev/mst197.

57. Lee J-Y, Baum SF, Oh S-H, Jiang C-Z, Chen J-C, Bowman JL. Recruitment of CRABS CLAW to promote nectary development within the eudicot clade. Development. 2005;132:5021–32. doi:10.1242/dev.02067.

58. Kram BW, Xu WW, Carter CJ. Uncovering the *Arabidopsis thaliana* nectary transcriptome: Investigation of differential gene expression in floral nectariferous tissues. BMC Plant Biol. 2009;9:92. doi:10.1186/1471-2229-9-92.

59. Thomas JB, Hampton ME, Dorn KM, David Marks M, Carter CJ. The pennycress (*Thlaspi arvense* L.) nectary: structural and transcriptomic characterization. BMC Plant Biol. 2017;17:201. doi:10.1186/s12870-017-1146-8.

60. Liu H, Ma J, Li H. Transcriptomic and microstructural analyses in *Liriodendron tulipifera* Linn. reveal candidate genes involved in nectary development and nectar secretion. BMC Plant Biol. 2019;19:531. doi:10.1186/s12870-019-2140-0.

61. Chatt EC, Mahalim S-N, Mohd-Fadzil N-A, Roy R, Klinkenberg PM, Horner HT, et al. Nectar biosynthesis is conserved among floral and extrafloral nectaries. Plant Physiol. 2021;185:1595–616. doi:10.1093/plphys/kiab018.

62. Gao X, Wang L, Zhang H, Zhu B, Lv G, Xiao J. Transcriptome analysis and identification of genes associated with floral transition and fruit development in rabbiteye blueberry (*Vaccinium ashei*). PLoS ONE. 2021;16:e0259119. doi:10.1371/journal.pone.0259119.

63. Devi A, Seth R, Masand M, Singh G, Holkar A, Sharma S, et al. Spatial genomic resource reveals molecular insights into key bioactive-metabolite biosynthesis in endangered *Angelica glauca* Edgew. Int J Mol Sci 2022. doi:10.3390/ijms231911064.

64. Zhang L, Wang L, Yang Y, Cui J, Chang F, Wang Y, Ma H. Analysis of *Arabidopsis* floral transcriptome: detection of new florally expressed genes and expansion of Brassicaceae-specific gene families. Front Plant Sci. 2014;5:802. doi:10.3389/fpls.2014.00802.

65. Milne RJ, Grof CP, Patrick JW. Mechanisms of phloem unloading: shaped by cellular pathways, their conductances and sink function. Curr Opin Plant Biol. 2018;43:8–15. doi:10.1016/j.pbi.2017.11.003.

66. Chen L-Q. SWEET sugar transporters for phloem transport and pathogen nutrition. New Phytol. 2014;201:1150–5. doi:10.1111/nph.12445.

67. Riens B, Lohaus G, Winter H, Heldt H. Production and diurnal utilization of assimilates in leaves of spinach (*Spinacia oleracea* L.) and barley (*Hordeum vulgare* L.). Planta. 1994;192:497–501. doi:10.1007/BF00203587.

68. Ma X-L, Milne RI, Zhou H-X, Song Y-Q, Fang J-Y, Zha H-G. Proteomics and post-secretory content adjustment of *Nicotiana tabacum* nectar. Planta. 2019;250:1703–15. doi:10.1007/s00425-019-03258-4.

69. Persia D, Cai G, Del Casino C, Faleri C, Willemse MTM, Cresti M. Sucrose synthase is associated with the cell wall of tobacco pollen tubes. Plant Physiol. 2008;147:1603–18. doi:10.1104/pp.108.115956.

70. Solhaug EM, Roy R, Chatt EC, Klinkenberg PM, Mohd-Fadzil N-A, Hampton M, et al. An integrated transcriptomics and metabolomics analysis of the *Cucurbita pepo* nectary implicates key modules of primary metabolism involved in nectar synthesis and secretion. Plant Direct. 2019b;3:e00120. doi:10.1002/pld3.120.

71. Ge YX, Angenent GC, Wittich PE, Peters J, Franken J, Busscher M, et al. NEC1, a novel gene, highly expressed in nectary tissue of *Petunia hybrida*. Plant J. 2000;24:725–34.

72. Schmölzer K, Gutmann A, Diricks M, Desmet T, Nidetzky B. Sucrose synthase: A unique glycosyltransferase for biocatalytic glycosylation process development. Biotechnol Adv. 2016;34:88–111. doi:10.1016/j.biotechadv.2015.11.003.

73. Eom J-S, Chen L-Q, Sosso D, Julius BT, Lin IW, Qu X-Q, et al. SWEETs, transporters for intracellular and intercellular sugar translocation. Curr Opin Plant Biol. 2015;25:53–62. doi:10.1016/j.pbi.2015.04.005.

74. Lin W, Pu Y, Liu S, Wu Q, Yao Y, Yang Y, et al. Genome-wide identification and expression patterns of AcSWEET family in pineapple and AcSWEET11 mediated sugar accumulation. Int J Mol Sci. 2022;23(22):13875. doi:10.3390/ijms232213875.

75. Denisow B, Masierowska M, Antoń S. Floral nectar production and carbohydrate composition and the structure of receptacular nectaries in the invasive plant *Bunias orientalis* L. (Brassicaceae). Protoplasma. 2016;253:1489–501. doi:10.1007/s00709-015-0902-6.

76. Carpaneto A, Geiger D, Bamberg E, Sauer N, Fromm J, Hedrich R. Phloem-localized, proton-coupled sucrose carrier ZmSUT1 mediates sucrose efflux under the control of the sucrose gradient and the proton motive force. J Biol Chem. 2005;280:21437–43. doi:10.1074/jbc.M501785200.

77. Zha H-G, Flowers VL, Yang M, Chen L-Y, Sun H. Acidic α-galactosidase is the most abundant nectarin in floral nectar of common tobacco (*Nicotiana tabacum*). Ann Bot. 2012;109:735–45. doi:10.1093/aob/mcr321.

78. Baker HG, Baker I. A brief historical review of the chemistry of floral nectar. Pp. 126-152. In: Bentley, E, editor. The biology of nectaries. Columbia University Press. 1983a. p.126–152.

79. Wist TJ, Davis AR. Floral nectar production and nectary anatomy and ultrastructure of *Echinacea purpurea* (Asteraceae). Ann Bot. 2006;97:177–93. doi:10.1093/aob/mcj027.

