## Supplementary Material for "Metabolic and transcriptomic analyses of nectaries reveal differences in the mechanism of nectar production between monocots (*Ananas comosus*) and dicots (*Nicotiana tabacum*)"

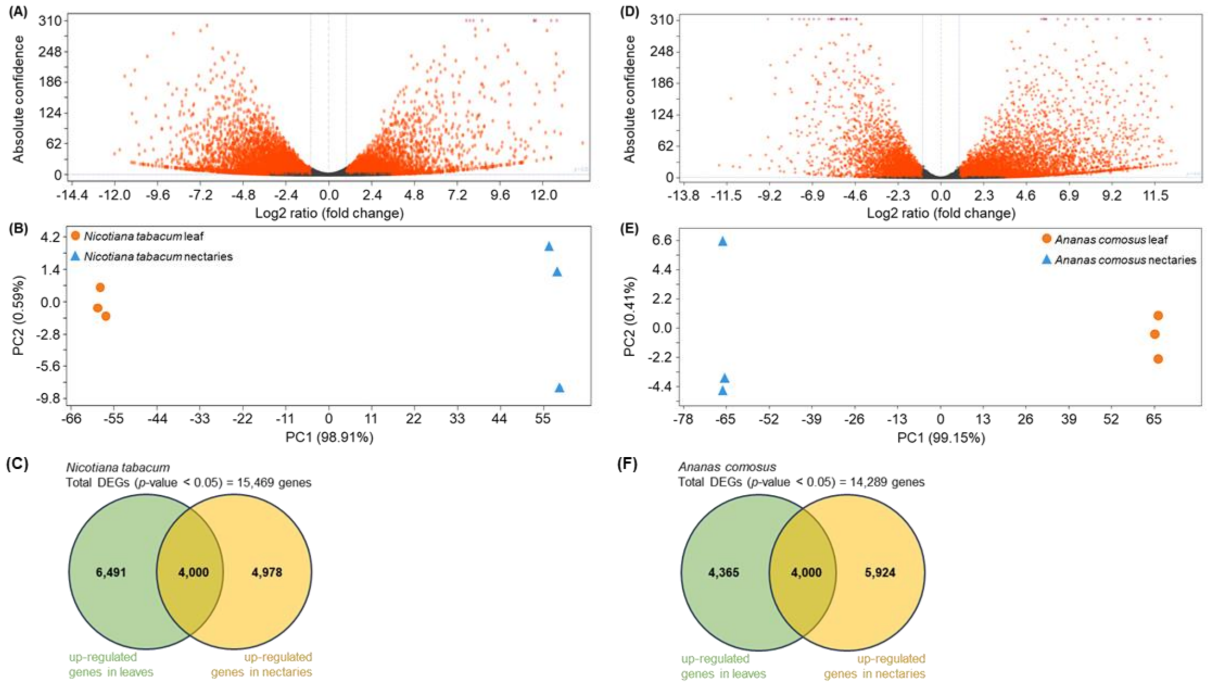


**Supplementary Figure S1**: **Volcano plot, PCA plot, and Venn diagram of the transcriptomic data of *N. tabacum* and *A. comosus*.**
The Volcano plot (A: *N. tabacum*, D: *A. comosus*) is generated by using expression levels using DESeq2 and shows the most highly differentially expressed loci. In this graphical presentation, fold change (log2 ratio) is plotted against absolute confidence (-log10 adjusted *p*-value). Each gene is represented by one dot and the orange dots have a *p*-value smaller than 0.05 in this plot and are the most differentially expressed genes. The red asterisks are genes with a *p*-value of zero. Principal component analysis (PCA) was used to visualize the variation between expression analysis samples (B: *N. tabacum*, E: *A. comosus*). The first principal component (PC 1) describes about 99 % and the second principal component (PC 2) describes less than 1 % of the dataset variation (B, E). In the Venn diagram, the DEGs (*p*-value < 0.05) are divided into up-regulated genes in leaves (green) and up-regulated genes in nectaries (yellow) for *Nicotiana tabacum* (C) and *Ananas comosus* (F). The number of genes in the overlapping of the two circles represents the number of genes that are neither up-regulated nor down-regulated and therefore cannot be clearly assigned to a tissue (C, F).

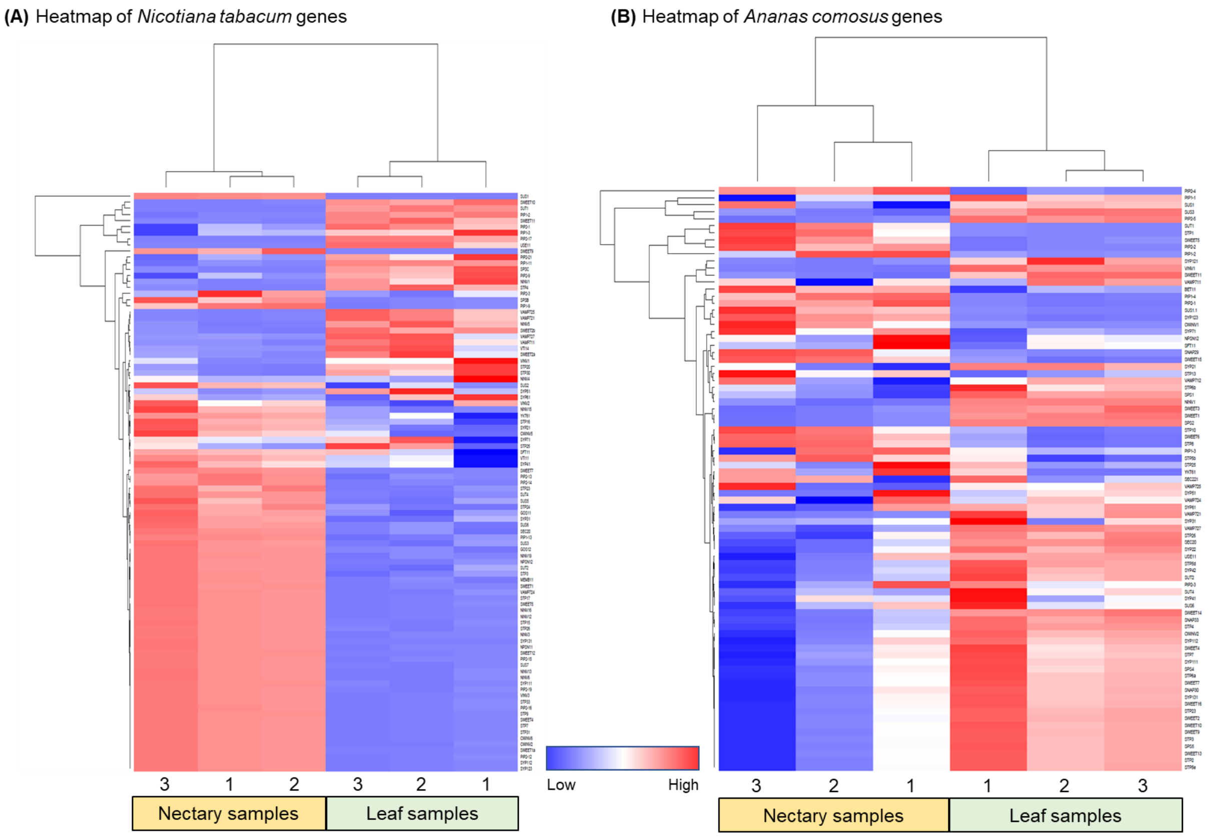


**Supplementary Figure S2: Expression heatmaps of the *Nicotiana tabacum* (A) and *Ananas comosus* (B) genes.**
The heatmap shows the scaled TPM values of different genes (SPS, INV, SUS, SUT, SWEET, STP, PIP, SNARE) of the individual samples of leaves and nectaries.

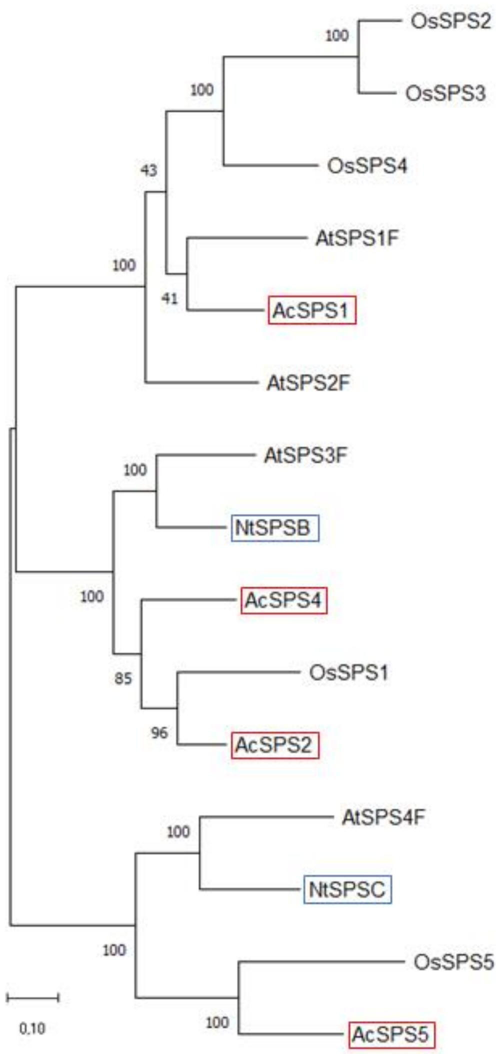


**Supplementary Figure S3**: **Phylogenetic analysis of selected sucrose phosphate synthases (SPS).** Phylogenetic analysis was carried out with the species *A. thaliana*, *N. tabacum*, *O. sativa*, and *A. comosus*. Protein alignment of the sucrose phosphate synthases was carried out by ClustalW. A maximum likelihood tree with 1,000 bootstrap iterations was calculated. Bar indicates evolutionary distance; numbers indicate percentage of bootstrap analysis. The red frame highlights the pineapple genes and the blue frame highlights the tobacco genes. Gene IDs are in Supplementary Table S4-S7.

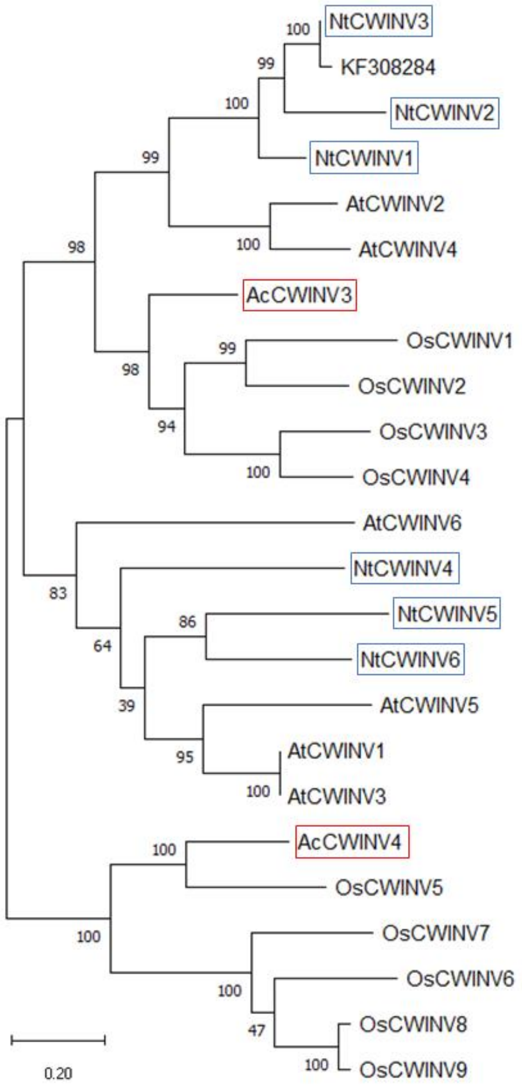


**Supplementary Figure S4**: **Phylogenetic analysis of selected cell wall invertases (CWINV).** Phylogenetic analysis was carried out with the species *A. thaliana*, *N. tabacum*, *O. sativa*, and *A. comosus*. Protein alignment of the invertases was carried out by ClustalW. A maximum likelihood tree with 1,000 bootstrap iterations was calculated. Bar indicates evolutionary distance; numbers indicate percentage of bootstrap analysis. The red frame highlights the pineapple genes and the blue frame highlights the tobacco genes. Gene IDs are in Supplementary Table S4-S7.

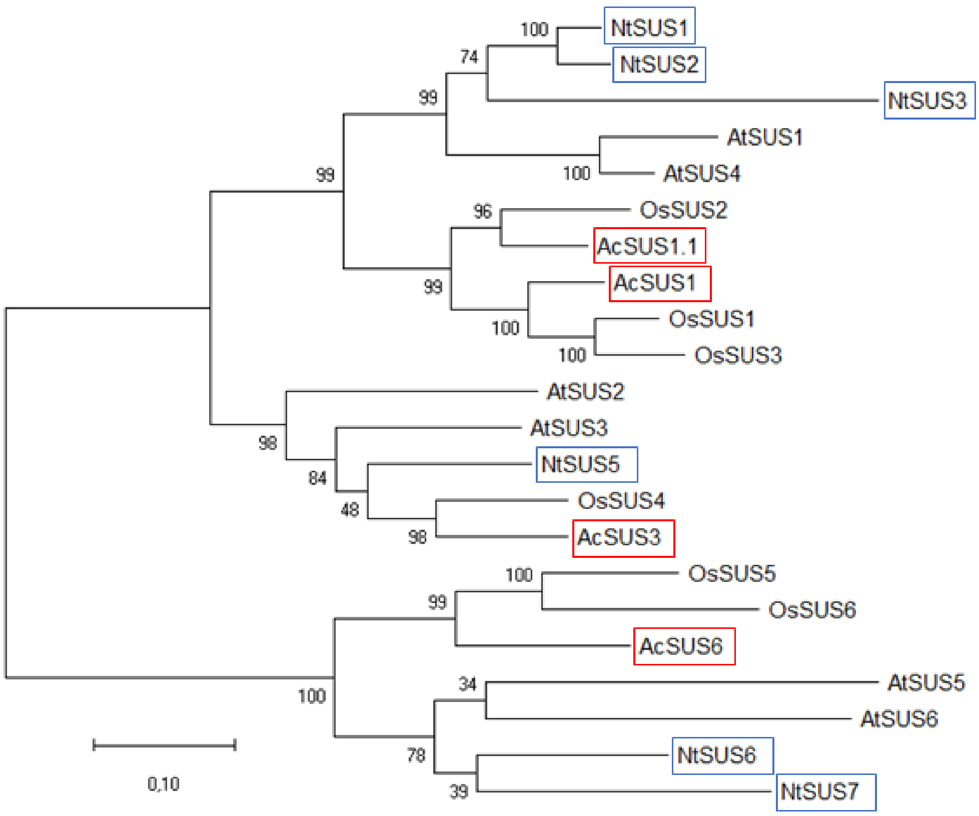


**Supplementary Figure S5**: **Phylogenetic analysis of selected sucrose synthases (SUS).** Phylogenetic analysis was carried out with the species *A. thaliana*, *N. tabacum*, *O. sativa*, and *A. comosus*. Protein alignment of the sucrose synthases was carried out by ClustalW. A maximum likelihood tree with 1,000 bootstrap iterations was calculated. Bar indicates evolutionary distance; numbers indicate percentage of bootstrap analysis. The red frame highlights the pineapple genes and the blue frame highlights the tobacco genes. Gene IDs are in Supplementary Table S4-S7.

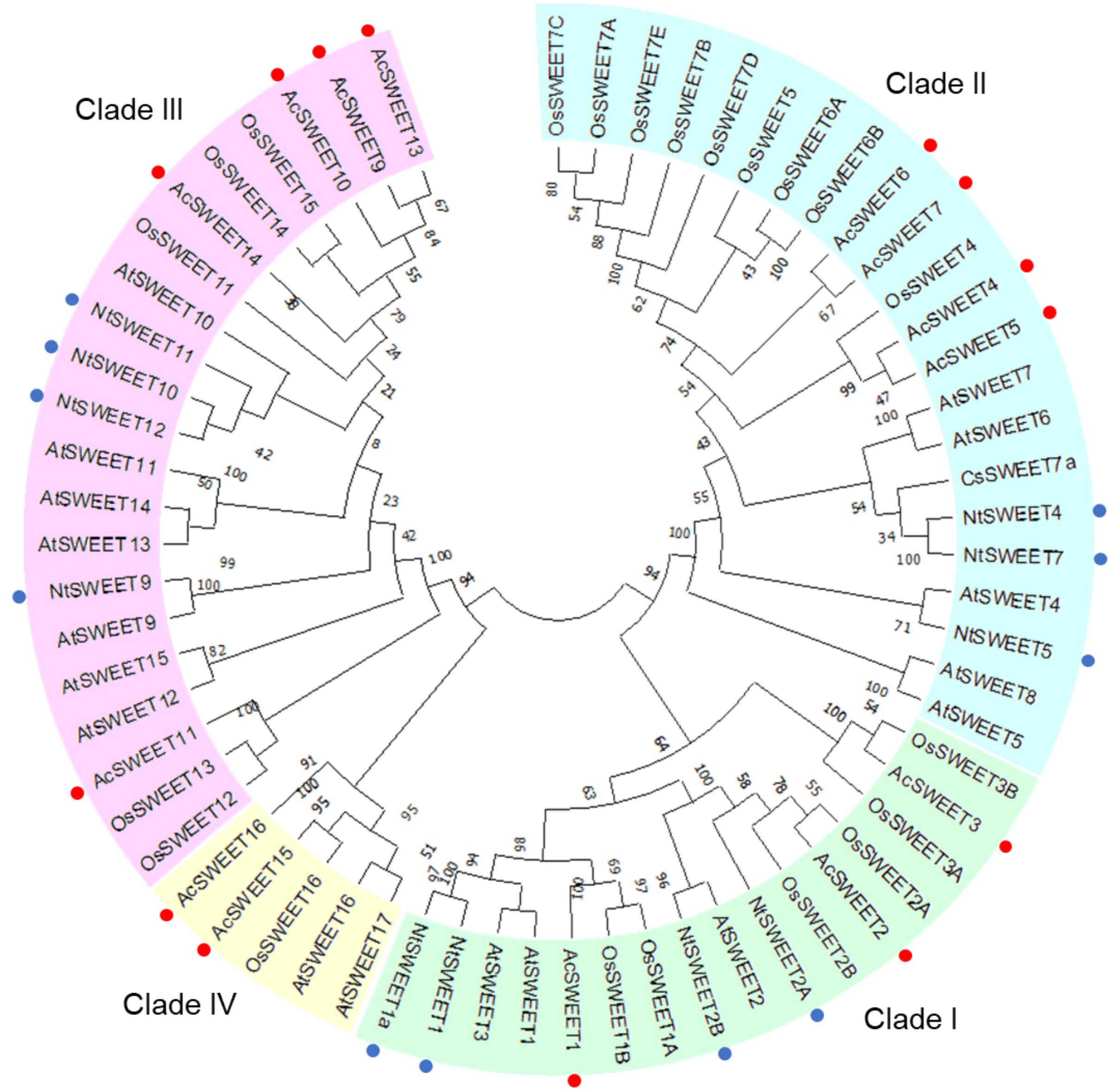


**Supplementary Figure S6**: **Phylogenetic analysis of selected sugars will eventually be exported transporters (SWEET).**

Phylogenetic analysis was carried out with the species *A. thaliana*, *N. tabacum*, *O. sativa*, and *A. comosus*. In addition, the sugar transporter CsSWEET7a was also used for this purpose, due to its role in the unloading of the apoplasmic phloem into nectar during cucumber anthesis (Li et al., 2021). Protein alignment of the transporters was carried out by ClustalW. A maximum likelihood tree with 1,000 bootstrap iterations was calculated. Bar indicates evolutionary distance; numbers indicate percentage of bootstrap analysis. SWEETs are divided into different clades depending on Lin *et al.*, 2022. The red point highlights the pineapple genes and the blue point highlights the tobacco genes. Gene IDs are in Supplementary Table S4-S7.

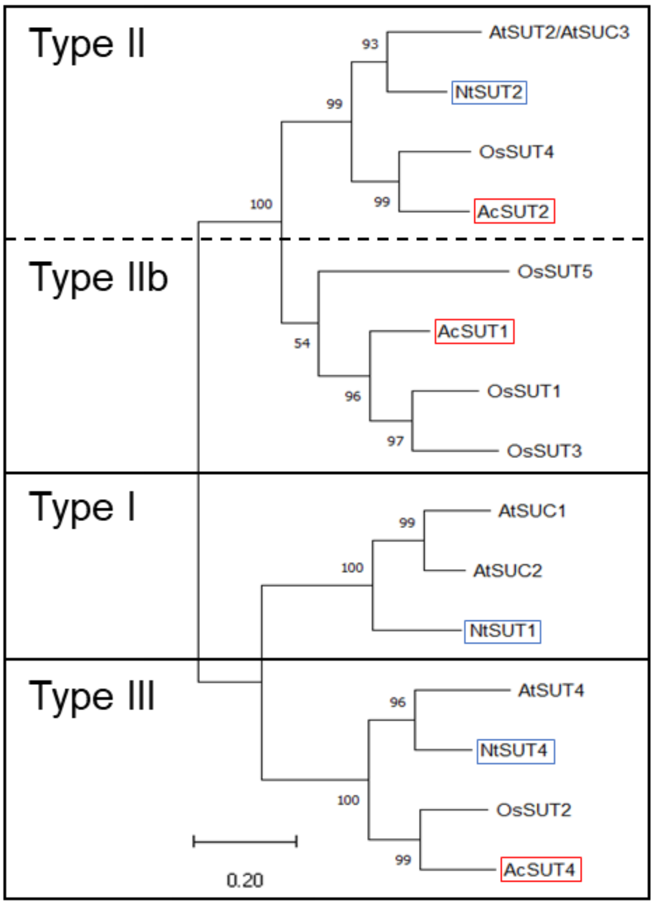


**Supplementary Figure S7**: **Phylogenetic analysis of selected sucrose uptake transporters (SUT).** Phylogenetic analysis was carried out with the species *A. thaliana*, *N. tabacum*, *O. sativa*, and *A. comosus*. Protein alignment of the sucrose transporter was carried out by ClustalW. A maximum likelihood tree with 1,000 bootstrap iterations was calculated. Bar indicates evolutionary distance; numbers indicate percentage of bootstrap analysis. SUTs are divided into different clades or types depending on Sauer 2007 and Peng et al. 2014. The red frame highlights the pineapple genes and the blue frame highlights the tobacco genes. Gene IDs are in Supplementary Table S4-S7.

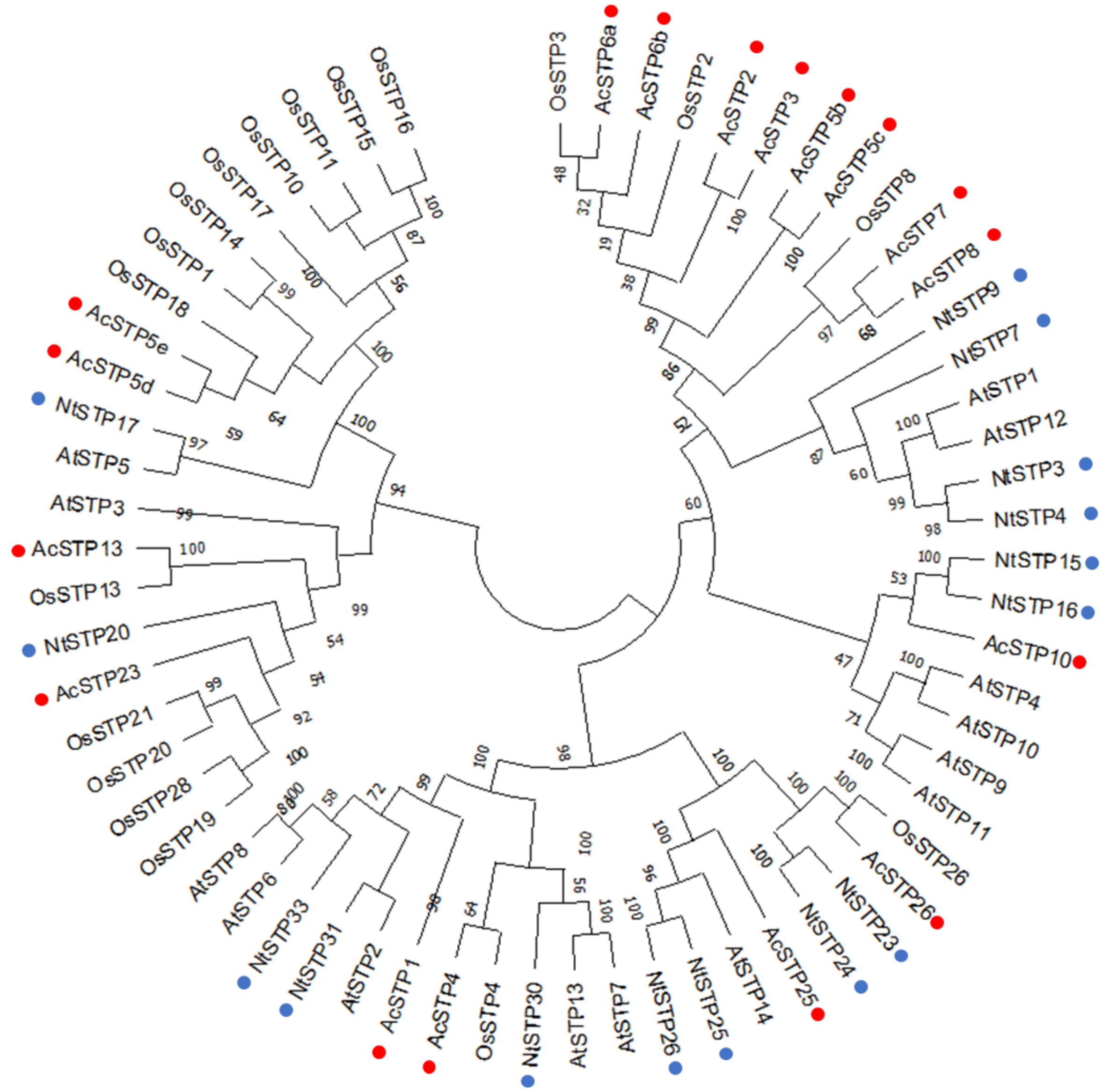


**Supplementary Figure S8**: **Phylogenetic analysis of selected sugar transport proteins (STP).** Phylogenetic analysis was carried out with the species *A. thaliana*, *N. tabacum*, *O. sativa*, and *A. comosus*. Protein alignment of the transport proteins was carried out by ClustalW. A maximum likelihood tree with 1,000 bootstrap iterations was calculated. Bar indicates evolutionary distance; numbers indicate percentage of bootstrap analysis. The red point highlights the pineapple genes and the blue point highlights the tobacco genes. Gene IDs are in Supplementary Table S4-S7.

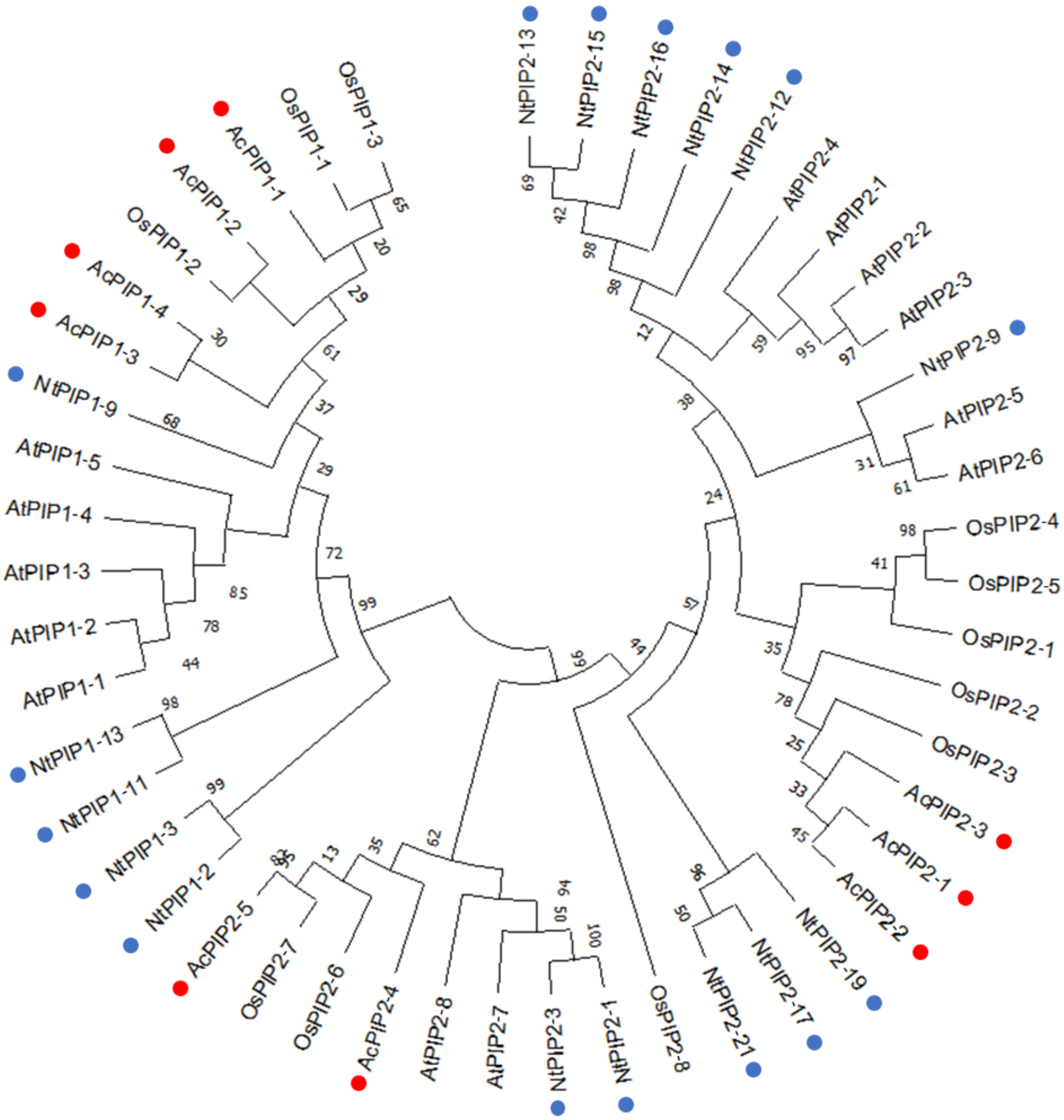


**Supplementary Figure S9**: **Phylogenetic analysis of selected plasma membrane intrinsic proteins (PIP).** Phylogenetic analysis was carried out with the species *A. thaliana*, *N. tabacum*, *O. sativa*, and *A. comosus*. Protein alignment of the intrinsic proteins was carried out by ClustalW. A maximum likelihood tree with 1,000 bootstrap iterations was calculated. Bar indicates evolutionary distance; numbers indicate percentage of bootstrap analysis. The red point highlights the pineapple genes and the blue point highlights the tobacco genes. Accession numbers are in Supplementary Table S4-S7.

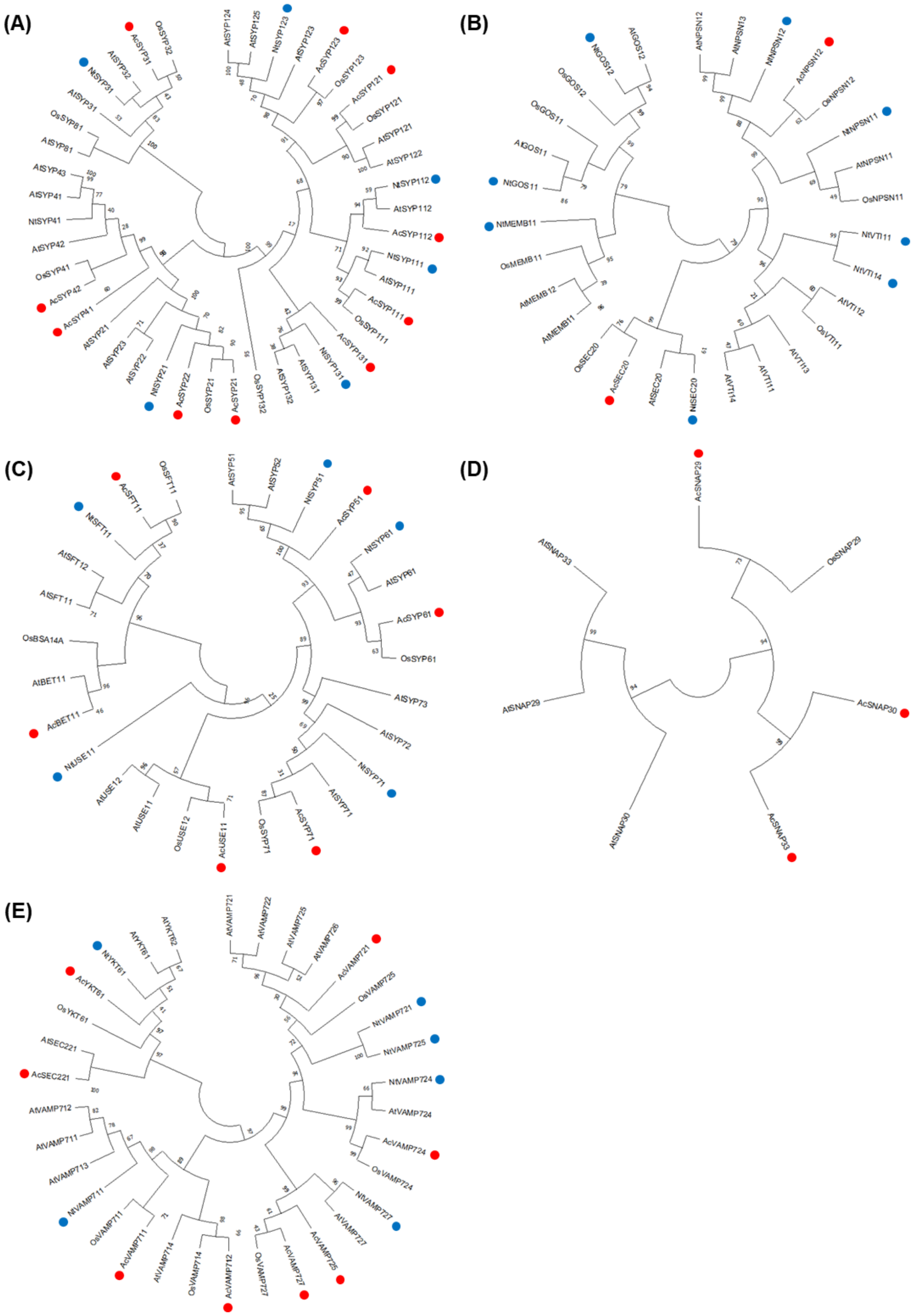


**Supplementary Figure S10**: **Phylogenetic analysis of selected soluble *N*-ethylmaleimide-sensitive-factor attachment receptor (SNARE)-domain-containing proteins.**
Phylogenetic analysis was carried out with the species *A. thaliana*, *N. tabacum*, *O. sativa*, and *A. comosus*. A separate phylogenetic tree was created for each clade (A: Clade Qa; B: Clade Qb; C: Clade Qc; D: Clade Qb + Qc; E: Clade R). Protein alignment of the proteins was carried out by ClustalW. A maximum likelihood tree with 1,000 bootstrap iterations was calculated. Bar indicates evolutionary distance; numbers indicate percentage of bootstrap analysis. The red point highlights the pineapple genes and the blue point highlights the tobacco genes. Gene IDs are in Supplementary Table S4-S7.

**Supplementary Table S1: Water content in nectaries and leaves, expressed as a percentage of fresh weight.**

| Species | Nectaries [%] | Leaves [%] |
|---|---|---|
| *Nicotiana tabacum* | 75.4 ± 1.7 | 85.6 ± 1.6 |
| *Ananas comosus* | 75.1 ±1.5 | 80.1 ± 1.7 |

**Supplementary Table S2: Differential expression log2 ratios of different genes in *Nicotiana tabacum*.**

The significant difference is indicated by asterisks (differential expression *p*-value < 0.05).

| Genes | Log2 ratio | Genes | Log2 ratio | Genes | Log2 ratio |
|---|---|---|---|---|---|
| *SPSB* | 6.4* | *SWEET10* | -1.5* | *PIP2-15* | 3.5* |
| *SPSC* | -2.3* | *SWEET11* | -4.3* | *PIP2-16* | 5.0* |
| *NINV1* | -0.8* | *SWEET12* | 4.0* | *PIP2-17* | -1.8* |
| *NINV3* | 1.6* | *SUT1* | -0.9* | *PIP2-19* | 0.0 |
| *NINV4* | 0.3 | *SUT2* | -0.8* | *PIP2-21* | 1.6* |
| *NINV5* | 0.6* | *SUT4* | 2.8* | *SYP111* | 2.2* |
| *NINV6* | 3.1* | *STP3* | -2.6* | *SYP112* | 1.1 |
| *NINV12* | -3.7* | *STP4* | -2.4* | *SYP123* | -0.5 |
| *NINV13* | 3.1* | *STP7* | -0.7 | *SYP131* | -0.2 |
| *NINV15* | 1.2* | *STP9* | 4.5* | *SYP21* | 0.2 |
| *NINV16* | -3.7* | *STP15* | 0.3 | *SYP31* | 0.9* |
| *NINV19* | 1.0* | *STP16* | -3.0* | *SYP41* | 0.8* |
| *VINV1* | 1.1* | *STP17* | -3.5* | *VTI11* | 1.2* |
| *VINV2* | 1.4* | *STP20* | -5.7* | *VTI14* | 0.5 |
| *VINV3* | -1.0* | *STP23* | 3.4* | *GOS11* | 0.1 |
| *CWINV2* | -0.3 | *STP24* | 2.3* | *GOS12* | 1.2* |
| *CWINV5* | -6.3* | *STP25* | -2.5* | *MEMB11* | 0.6* |
| *CWINV6* | 2.8 | *STP26* | -3.1* | *NPSN11* | -0.8* |
| *SUS1* | 8.6* | *STP30* | -4.2* | *NPSN12* | 0.4* |
| *SUS2* | 2.3* | *STP31* | -0.3 | *SEC20* | 1.0* |
| *SUS3* | -0.2 | *STP33* | -0.5 | *SYP51* | 0.0 |
| *SUS5* | 2.4* | *PIP1-2* | 0.2 | *SYP61* | -0.6* |
| *SUS6* | 0.5* | *PIP1-3* | 2.3* | *SYP71* | -0.5 |
| *SUS7* | 2.5* | *PIP1-9* | 4.6* | *SFT11* | 0.7 |
| *SWEET1* | 1.5* | *PIP1-11* | 0.6* | *USE11* | -0.4 |
| *SWEET1a* | -0.2 | *PIP1-13* | 1.0* | *VAMP711* | 0.5 |
| *SWEET2a* | -0.2 | *PIP2-1* | 2.3* | *VAMP721* | 0.3 |
| *SWEET2b* | -1.4* | *PIP2-3* | 3.6* | *VAMP724* | 0.6* |
| *SWEET4* | 8.3* | *PIP2-9* | 1.5* | *VAMP725* | 0.3 |
| *SWEET5* | -0.6 | *PIP2-12* | 1.8* | *VAMP727* | 0.4 |
| *SWEET7* | 8.9* | *PIP2-13* | 4.1* | *YKT61* | 1.0* |
| *SWEET9* | 12.8* | *PIP2-14* | 4.8* | | |

**Supplementary Table S3: Differential expression log2 ratios of different genes in *Ananas comosus.***

The significant difference is indicated by asterisks (differential expression *p*-value < 0.05).

| Genes | Log2 ratio | Genes | Log2 ratio | Genes | Log2 ratio |
|---|---|---|---|---|---|
| SPS1 | -1.2* | SUT4 | 1.4* | SYP121 | -2.9* |
| SPS2 | -4.3* | STP1 | 9.3* | SYP123 | 9.9* |
| SPS4 | 5.1* | STP2 | -0.5 | SYP131 | 3.6* |
| SPS5 | 0.6* | STP3 | 0.4 | SYP21 | -0.8* |
| NINV1 | -1.6* | STP4 | -4.3* | SYP22 | -0.3 |
| VINV1 | -3.6* | STP5b | 6.9* | SYP31 | 1.0* |
| CWINV1 | 3.2* | STP5d | -0.6* | SYP41 | 4.3* |
| CWINV2 | -0.4 | STP5e | -0.9 | SYP42 | 0.3 |
| SUS1 | -0.7* | STP6a | 5.0* | NPSN12 | 0.4 |
| SUS1.1 | 4.6* | STP6b | -0.9* | SEC20 | -0.1 |
| SUS3 | 4.6* | STP7 | 9.6* | BET11 | 0.5* |
| SUS6 | -2.6* | STP8 | 5.6* | SYP51 | 0.5* |
| SWEET1 | -4.4* | STP10 | 8.4* | SYP61 | 0.5* |
| SWEET2 | -0.6* | STP13 | 0.4* | SYP71 | 0.6* |
| SWEET3 | -5.2* | STP23 | -0.3 | SFT11 | 0.6 |
| SWEET4 | 2.2* | STP25 | 2.7* | USE11 | 0.3 |
| SWEET5 | 1.6* | STP26 | -0.1 | SNAP29 | 1.2* |
| SWEET6 | 8.7* | PIP1-1 | -1.3* | SNAP30 | 1.6* |
| SWEET7 | 5.3* | PIP1-2 | 2.8* | SNAP33 | -2.8* |
| SWEET9 | 0.3 | PIP1-3 | 2.2* | VAMP711 | -0.5 |
| SWEET10 | 4.5* | PIP1-4 | 2.7* | VAMP712 | -0.2 |
| SWEET11 | -2.1* | PIP2-1 | 3.9* | VAMP721 | 0.0 |
| SWEET13 | 0.4* | PIP2-2 | 2.0* | VAMP724 | 0.4* |
| SWEET14 | -3.1* | PIP2-3 | 3.4* | VAMP725 | 0.3 |
| SWEET15 | 7.5* | PIP2-4 | 1.1* | VAMP727 | -0.6* |
| SWEET16 | 2.6* | PIP2-5 | -3.6* | YKT61 | 0.8* |
| SUT1 | 3.6* | SYP111 | 2.8* | SEC221 | 0.7* |
| SUT2 | -0.3 | SYP112 | 1.3* | | |

**Supplementary Table S4: Gene IDs of *Arabidopsis thaliana* genes.** Genes are divided into the following groups: SPS, INV, SUS, SUT, SWEET, STP, PIP, SNARE. The genes can be found with the gene ID in the NCBI database.

| Gene common name | Gene ID | Gene common name | Gene ID | Gene common name | Gene ID |
|---|---|---|---|---|---|
| *AtSPS1F* | At5g20280 | *AtSTP3* | AT5G61520 | *AtSYP131* | AT3G03800 |
| *AtSPS2F* | At5g11110 | *AtSTP4* | AT3G19930 | *AtSYP132* | AT5G08080 |
| *AtSPS3F* | At1g04920 | *AtSTP5* | AT1G34580 | *AtSYP21* | AT5G16830 |
| *AtSPS4F* | At4g10120 | *AtSTP6* | AT3G05960 | *AtSYP22* | AT5G46860 |
| *AtNINV1* | At1g56560 | *AtSTP7* | AT4G02050 | *AtSYP23* | AT4G17730 |
| *AtCWINV1* | At3g13790 | *AtSTP8* | AT5G26250 | *AtSYP31* | AT5G05760 |
| *AtCWINV2* | At3g52600 | *AtSTP9* | AT1G50310 | *AtSYP32* | AT3G24350a |
| *AtCWINV4* | At2g36190 | *AtSTP10* | AT3G19940 | *AtSYP41* | AT5G26980 |
| *AtCWINV5* | At3g13784 | *AtSTP11* | AT5G23270 | *AtSYP42* | AT4G02195 |
| *AtCWINV6* | At5g11920 | *AtSTP12* | AT4G21480 | *AtSYP43* | AT3G05710 |
| *AtSus1* | At5g20830 | *AtSTP13* | AT5G26340 | *AtSYP81* | AT1G51740 |
| *AtSus2* | At5g49190 | *AtSTP14* | AT1G77210 | *AtVTI11* | AT5G39510 |
| *AtSus3* | At4g02280 | *AtSUC1* | AEE35247.1 | *AtVTI12* | AT1G26670 |
| *AtSus4* | At3g43190 | *AtSUC2* | AEC05635.1 | *AtVTI13* | AT3G29100 |
| *AtSus5* | At5g37180 | *AtSUT4* | NP_172467.1 | *AtVTI14* | AT5G39630 |
| *AtSus6* | At1g73370 | *AtPIP1-1* | AT3G61430 | *AtGOS11* | AT1G15880 |
| *AtSWEET1* | At1g21460 | *AtPIP1-2* | AT2G45960 | *AtGOS12* | AT2G45200 |
| *AtSWEET2* | At3g14770 | *AtPIP1-3* | AT1G01620 | *AtMEMB11* | AT2G36900 |
| *AtSWEET3* | At5g53190 | *AtPIP1-4* | AT4G00430 | *AtMEMB12* | AT5G50440 |
| *AtSWEET4* | At3g28007 | *AtPIP1-5* | AT4G23400 | *AtNPSN11* | AT2G35190 |
| *AtSWEET5* | At5g62850 | *AtPIP2-1* | AT3G53420 | *AtNPSN12* | AT1G48240 |
| *AtSWEET6* | At1g66770 | *AtPIP2-2* | AT2G37170 | *AtNPSN13* | AT3G17440 |
| *AtSWEET7* | At4g10850 | *AtPIP2-3* | AT2G37180 | *AtSEC20* | AT3G24315 |
| *AtSWEET8* | At5g40260 | *AtPIP2-4* | AT5G60660 | *AtBET11* | AT3G58170 |
| *AtSWEET9* | At2g39060 | *AtPIP2-5* | AT3G54820 | *AtSYP51* | AT1G16240 |
| *AtSWEET10* | At5g50790 | *AtPIP2-6* | AT2G39010 | *AtSYP52* | AT1G79590 |
| *AtSWEET11* | At3g48740 | *AtPIP2-7* | AT4G35100 | *AtSYP61* | AT1G28490 |
| *AtSWEET12* | At5g23660 | *AtPIP2-8* | AT2G16850 | *AtSYP71* | AT3G09740 |
| *AtSWEET13* | At5g50800 | *AtSYP111* | AT1G08560 | *AtSYP72* | AT3G45280 |
| *AtSWEET14* | At4g25010 | *AtSYP112* | AT2G18260 | *AtSYP73* | AT3G61450 |
| *AtSWEET15* | At5g13170 | *AtSYP121* | AT3G11820 | *AtSFT11* | AT4G14600 |
| *AtSWEET16* | At3g16690 | *AtSYP122* | AT3G52400 | *AtSFT12* | AT1G29060 |
| *AtSWEET17* | At4g15920 | *AtSYP123* | AT4G3330 | *AtUSE11* | AT1G54110 |
| *AtSTP1* | AT1G11260 | *AtSYP124* | AT1G61290 | *AtUSE12* | AT3G55600 |
| *AtSTP2* | AT1G07340 | *AtSYP125* | AT1G11250 | *AtSNAP29* | AT5G07880 |

| Gene common name | Gene ID |
|---|---|
| *AtSNAP30* | AT1G13890 |
| *AtSNAP33* | AT5G61210 |
| *AtVAMP711* | AT4G32150 |
| *AtVAMP712* | AT2G25340 |
| *AtVAMP713* | AT5G11150 |
| *AtVAMP714* | AT5G22360 |
| *AtVAMP721* | AT1G04750 |
| *AtVAMP722* | AT2G33120 |
| *AtVAMP724* | AT4G15780 |
| *AtVAMP725* | AT2G32670 |
| *AtVAMP726* | AT1G04760 |
| *AtVAMP727* | AT3G54300 |
| *AtYKT61* | AT5G58060 |
| *AtYKT62* | AT5G58180 |
| *SEC221* | AT1G11890 |

111

**Supplementary Table S5: Gene IDs of *Nicotiana tabacum* genes.**

Genes are divided into the following groups: SPS, INV, SUS, SUT, SWEET, STP, PIP, SNARE. The genes can be found with the gene ID in the NCBI database and with another gene ID in the *N. tabacum* genome (Edwards et al., 2017). The genes were named using the following references: Chen et al., 2005; Wang et al., 2015; Pfister et al, 2017; Ahmed et al., 2020; Xu et al., 2022; Cheng et al., 2023.

| Gene common name | Gene ID | Gene ID Genome |
|---|---|---|
| *NtSPSB* | DQ213015 | 0000401g0120 |
| *NtSPSC* | DQ213014 | 0000009g0480 |
| *NtNINV1* | LOC107778980 | 0003838g0090 |
| *NtNINV3* | LOC107819026 | 0001082g0080 |
| *NtNINV4* | LOC107816919 | 0000962g0090 |
| *NtNINV5* | LOC107768486 | 0002561g0010 |
| *NtNINV6* | LOC107781195 | 0000589g0010 |
| *NtNINV8* | LOC107768486 | 0002561g0010 |
| *NtNINV12* | LOC107797521 | 0000283g0090 |
| *NtNINV13* | LOC107781195 | 0000589g0010 |
| *NtNINV14* | LOC107778980 | 0003838g0090 |
| *NtNINV15* | LOC107826453 | 0000365g0230 |
| *NtNINV16* | LOC107797521 | 0000283g0090 |
| *NtNINV19* | LOC107788023 | 0000303g0260 |
| *NtNINV20* | LOC107768486 | 0002561g0010 |
| *NtCWINV2* | AF376773.1 | 0011403g0010 |
| *NtCWINV5* | HM022265.1 | 0001295g0210 |
| *NtCWINV6* | ADI70683.1 | 0002654g0150 |
| *NtVINV1* | LOC107770131 | 0001383g0030 |
| *NtVINV2* | LOC107810155 | 0001780g0120 |
| *NtVINV3* | LOC107807993 | 0000993g0100 |
| *NtSus1* | LOC107783276 | 0000073g0400 |
| *NtSus2* | LOC107775654 | 0002280g0010 |
| *NtSus3* | LOC107804066 | 0000170g0050 |
| *NtSus5* | LOC107775584 | 0000116g0360 |
| *NtSus6* | LOC107766404 | 0000483g0250 |
| *NtSus7* | LOC107789475 | 0001180g0180 |
| *NtSWEET1* | LOC107806439 | 0000445g0160.1 |
| *NtSWEET1a* | LOC107787259 | 0004095g0060.1 |
| *NtSWEET2A* | XM_016629722.1 | 0003746g0030 |
| *NtSWEET2B* | XM_016634622.1 | 0001140g0150 |
| *NtSWEET4* | LOC107804242 | 0000673g0010.1 |
| *NtSWEET5* | LOC107797107 | 0000003g0510.1 |
| *NtSWEET7* | LOC107794016 | 0002367g0130.1 |
| *NtSWEET9* | LOC107793141 | 0004802g0020 |
| *NtSWEET10* | LOC107788219 | 0001332g0120.1 |
| *NtSWEET11* | LOC107760004 | 0000033g0010.1 |
| *NtSWEET12* | LOC107760005 | 0000033g0030.1 |
| *NtSTP3* | XP_016474706.1 | 0003465g0030 |
| *NtSTP4* | XP_016494599.1 | 0000190g0030 |
| *NtSTP7* | XP_016476713.1 | 0002234g0120 |
| *NtSTP9* | XP_016484433.1 | 0004425g0040 |
| *NtSTP15* | XP_016475527.1 | 0002387g0060 |
| *NtSTP16* | XP_016488213.1 | 0000069g0190 |
| *NtSTP17* | XP_016479727.1 | 0000232g0280 |
| *NtSTP20* | XP_016434630.1 | 0001267g0050 |
| *NtSTP23* | XP_016435043.1 | 0000416g0010 |
| *NtSTP24* | XP_016493725.1 | 0001671g0060 |
| *NtSTP25* | XP_016446212.1 | 0000012g0180 |
| *NtSTP26* | XP_016483827.1 | 0001004g0170 |
| *NtSTP30* | XP_016500201.1 | 0000795g0070 |
| *NtSTP31* | XP_016458581.1 | 0000102g0190 |
| *NtSTP33* | XP_016515629.1 | 0000559g0110 |
| *NtSUT1* | MF140390.1 | 0000159g0100 |
| *NtSUT2* | LC497468.1 | 0003062g0010 |
| *NtSUT4* | AB539539.1 | 0000377g0120 |
| *NtPIP1-2* | XP_016508253.1 | 0001615g0140 |
| *NtPIP1-3* | AAB04757.1 | 0003043g0010 |
| *NtPIP1-9* | NP_001312921.1 | 0000737g0120 |
| *NtPIP1-11* | XP_016515710.1 | 0000846g0060 |
| *NtPIP1-13* | XP_016510215.1 | 0000583g0150 |
| *NtPIP2-1* | AAL33586.1 | 0000283g0420 |
| *NtPIP2-3* | NP_001312414.1 | 0003914g0040 |
| *NtPIP2-9* | NP_001312511.1 | 0000575g0130 |
| *NtPIP2-12* | NP_001312276.1 | 0001192g0080 |
| *NtPIP2-13* | NP_001312334.1 | 0009795g0010 |
| *NtPIP2-14* | XP_016486700.1 | 0000101g0120 |
| *NtPIP2-15* | NP_001312333.1 | 0009795g0020 |

| Gene common name | Gene ID | Gene ID Genome |
|---|---|---|
| *NtPIP2-16* | XP_016513533.1 | 0000101g0110 |
| *NtPIP2-17* | NP_001312464.1 | 0000650g0260 |
| *NtPIP2-19* | NP_001313208.1 | 0000106g0170 |
| *NtPIP2-21* | NP_001311765.1 | 0000181g0120 |
| *NtSYP111* | XP_016479641.1 | 0000302g0040 |
| *NtSYP112* | XP_016489059.1 | 0000178g0230 |
| *NtSYP123* | XP_016462928.1 | 0000635g0160 |
| *NtSYP131* | XP_016510458.1 | 0004274g0010 |
| *NtSYP21* | XP_016506496.1 | 0000109g0090 |
| *NtSYP31* | XP_016462826.1 | 0000125g0090 |
| *NtSYP41* | XP_016443453.1 | 0002715g0150 |
| *NtVTI11* | XP_016500539.1 | 0001997g0110 |
| *NtVTI14* | NP_001312652.1 | 0001195g0020 |
| *NtGOS11* | XP_016486928.1 | 0002503g0100 |
| *NtGOS12* | XP_016482025.1 | 0000859g0340 |
| *NtMEMB11* | XP_016454786.1 | 0001741g0020 |
| *NtNPSN11* | XP_016440521.1 | 0001981g0010 |
| *NtNPSN12* | XP_016513453.1a | 0000895g0240 |
| *NtSEC20* | XP_016474547.1 | 0001041g0140 |
| *NtSYP51* | XP_016487954.1 | 0001160g0040 |
| *NtSYP61* | XP_016456479.1 | 0000151g0150 |
| *NtSYP71* | XP_016478104.1 | 0001192g0040 |
| *NtSFT11* | XP_016464993.1 | 0001984g0080 |
| *NtUSE11* | XP_016471202.1 | 0000630g0070 |
| *NtVAMP711* | XP_016451055.1 | 0000109g0410 |
| *NtVAMP721* | XP_016433615.1 | 0002085g0030 |
| *NtVAMP724* | XP_016443055.1 | 0000960g0120 |
| *NtVAMP725* | XP_016507558.1 | 0002085g0030 |
| *NtVAMP727* | XP_016457065.1 | 0002586g0070 |
| *NtYKT61* | XP_016496169.1 | 0001094g0120 |

**Supplementary Table S6: Gene IDs of *Oryza sativa* genes.**
Genes are divided into the following groups: SPS, INV, SUS, SUT, SWEET, STP, PIP, SNARE. The genes can be found with the gene ID in the NCBI database.

| Gene common name | Gene ID |
|---|---|
| *OsSPS1* | Q0JGK4 |
| *OsSPS2* | B7F7B9.2 |
| *OsSPS3* | Q67WN8 |
| *OsSPS4* | Q6ZHZ1 |
| *OsSPS5* | Q53JI9 |
| *OsNINV1* | Os02g34560 |
| *OsNINV2* | Os04g35280 |
| *OsNINV3* | Os11g07440 |
| *OsNINV4* | Os02g03320 |
| *OsNINV5* | Os02g32730 |
| *OsNINV6* | Os04g33490 |
| *OsNINV7* | Os01g22900 |
| *OsNINV8* | Os03g20020 |
| *OsCWINV1* | Os03g52560 |
| *OsCWINV2* | Os04g33720 |
| *OsCWINV3* | Os04g33740 |
| *OsCWINV4* | Os02g33110 |
| *OsCWINV5* | Os01g73580 |
| *OsCWINV6* | Os04g56930 |
| *OsCWINV7* | Os04g56920 |
| *OsCWINV8* | Os09g08120 |
| *OsCWINV9* | Os09g08072 |
| *OsVINV1* | Os02g01590 |
| *OsVINV2* | Os04g45290 |
| *OsSus1* | NP_001389102.1 |
| *OsSus2* | NP_001389706.1 |
| *OsSus3* | NP_001390086.1 |
| *OsSus4* | NP_001389082.1 |
| *OsSus5* | NP_001406164.1 |
| *OsSus6* | NP_001388945.1 |
| *OsSWEET1A* | AK099531 |
| *OsSWEET1B* | AK063475 |
| *OsSWEET2A* | AK104255 |
| *T2B* | AK059965 |
| *OsSWEET3A* | NP_001407211.1 |

| Gene common name | Gene ID |
|---|---|
| *OsSWEET3B* | XP_015642315.1 |
| *OsSWEET4* | AK071676 |
| *OsSWEET5* | AK069614 |
| *OsSWEET6A* | NP_001415004.1 |
| *OsSWEET6B* | AK099440 |
| *OsSWEET7A* | Q0J361.2 |
| *OsSWEET7B* | XP_015611083.1 |
| *OsSWEET7C* | XP_015619157.1 |
| *OsSWEET7D* | B9G2E6.3 |
| *OsSWEET7E* | A3BWJ9.2 |
| *OsSWEET11* | AK106127 |
| *OsSWEET12* | AK109114 |
| *OsSWEET13* | CI437556 |
| *OsSWEET14* | AK101913 |
| *OsSWEET15* | AK103266 |
| *OsSWEET16* | CI149956 |
| *OsSTP1* | Os04g37980.1 |
| *OsSTP2* | Os03g39710.1 |
| *OsSTP3* | Os07g01560.1 |
| *OsSTP4* | Os03g11900.1 |
| *OsSTP8* | Os01g38670.1 |
| *OsSTP10* | Os02g36414.1 |
| *OsSTP11* | Os02g36440.1 |
| *OsSTP13* | Os03g01170.1 |
| *OsSTP14* | Os04g37970.1 |
| *OsSTP15* | Os04g37990.1 |
| *OsSTP16* | Os04g38010.1 |
| *OsSTP17* | Os04g38026.1 |
| *OsSTP18* | Os04g38220.1 |
| *OsSTP19* | Os06g04900.1 |
| *OsSTP20* | Os07g03910.1 |
| *OsSTP21* | Os07g03960.1 |
| *OsSTP26* | Os09g24924.1 |
| *OsSTP28* | Os11g38160.1 |
| *OsSUT1* | BAA24071 |

| Gene common name | Gene ID |
|---|---|
| *OsSUT2* | AB091672 |
| *OsSUT3* | BAB68368 |
| *OsSUT4* | AB091673 |
| *OsSUT5* | AB091674 |
| *OsPIP1-1* | BAD28398 |
| *OsPIP1-2* | Os04g47220 |
| *OsPIP1-3* | BAD22920 |
| *OsPIP2-1* | BAC15868 |
| *OsPIP2-2* | BAD23735 |
| *OsPIP2-3* | CAD41442 |
| *OsPIP2-4* | BAC16113 |
| *OsPIP2-5* | BAC16116 |
| *OsPIP2-6* | CAE05002 |
| *OsPIP2-7* | BAD46581 |
| *OsPIP2-8* | AAP44741 |
| *OsSYP81* | XP_015635852.1 |
| *OsSYP32* | XP_015617756.1 |
| *OsSYP41* | XP_015641289.1 |
| *OsSYP21* | XP_015622380.1 |
| *OsSYP111* | XP_015631526.1 |
| *OsSYP121* | XP_015628438.1 |
| *OsSYP123* | XP_015624037.1 |
| *OsSYP132* | XP_015647004.1 |
| *OsSEC20* | XP_015623272.1 |
| *OsMemB11* | XP_015633085.1 |
| *OsGOS11* | XP_015611820.1 |
| *OsGOS12* | XP_015627527.1 |
| *OsVTI11* | XP_015621845.1 |
| *OsNPSN11* | XP_015642684.1 |
| *OsNPSN12* | XP_015646134.1 |
| *OsUSE12* | XP_015651428.1 |
| *OsBSA14A* | XP_015648254.1 |
| *OsSFT11* | XP_015645414.1 |
| *OsSYP61* | XP_015625156.1 |
| *OsSYP71* | XP_015639751.1 |

| **Gene common name** | **Gene ID** |
|---|---|
| *OsSNAP29* | NP_001403563.1 |
| *OSYKT61* | NP_001393116.1 |
| *OsVAMP711* | XP_015641118.1 |
| *OsVAMP714* | XP_015614986.1 |
| *OsVAMP724* | XP_015645337.1 |
| *OsVAMP727* | XP_015649386.1 |
| *OsVAMP725* | XP_015647709.1 |

**Supplementary Table S7: Gene IDs of *Ananas comosus* genes.**

Genes are divided into the following groups: SPS, INV, SUS, SUT, SWEET, STP, PIP, SNARE. The genes can be found with the gene ID in the NCBI database and in the *A. comosus* genome (Ming et al., 2015). The genes were named using the following references: Guo et al., 2018; Zhu and Ming, 2019; Wu et al., 2022; Gao et al., 2022; Fakher et al., 2022.

| **Gene common name** | **Gene ID** |
|---|---|
| *AcSPS1* | LOC109724677 |
| *AcSPS2* | LOC109712530 |
| *AcSPS4* | LOC109725728 |
| *AcSPS5* | LOC109726629 |
| *AcNINV1* | LOC109726732 |
| *AcNINV2* | LOC109707584 |
| *AcNINV3* | LOC109712448 |
| *AcNINV4* | LOC109728768 |
| *AcNINV5* | LOC109707134 |
| *AcCWINV1* | LOC109725952 |
| *AcCWINV2* | LOC109725753 |
| *AcVINV1* | LOC109727167 |
| *AcSus1* | LOC109714349 |
| *AcSus1.1* | LOC109717643 |
| *AcSus3* | LOC109726980 |
| *AcSus6* | LOC109719969 |
| *AcSWEET1* | LOC109717722 |
| *AcSWEET2* | LOC109717521 |
| *AcSWEET3* | LOC109716017 |
| *AcSWEET4* | LOC109710588 |
| *AcSWEET5* | LOC109717849 |
| *AcSWEET6* | LOC109716724 |
| *AcSWEET7* | LOC109722106 |
| *AcSWEET9* | LOC109723368 |
| *AcSWEET10* | LOC109725827 |
| *AcSWEET11* | LOC109723713 |
| *AcSWEET13* | LOC109726183 |
| *AcSWEET14* | LOC109710404 |
| *AcSWEET15* | LOC109709226 |
| *AcSWEET16* | LOC109720315 |
| *AcSTP1* | LOC109712863 |
| *AcSTP2* | LOC109708566 |
| *AcSTP3* | LOC109708611 |
| *AcSTP4* | LOC109721830 |
| *AcSTP5b* | LOC109718670 |
| *AcSTP5c* | LOC109718670 |
| *AcSTP5d* | LOC109725394 |
| *AcSTP5e* | LOC109725502 |
| *AcSTP6a* | LOC109713301 |
| *AcSTP6b* | LOC109715036 |
| *AcSTP7* | LOC109718125 |
| *AcSTP8* | LOC109724884 |
| *AcSTP10* | LOC109707304 |
| *AcSTP13* | LOC109713738 |
| *AcSTP23* | LOC109726972 |
| *AcSTP25* | LOC109708573 |
| *AcSTP26* | LOC109706529 |
| *AcSUT1* | LOC109720885 |
| *AcSUT2* | LOC109727628 |
| *AcSUT4* | LOC109718518 |
| *AcPIP1-1* | LOC109708083 |
| *AcPIP1-2* | LOC109703497 |
| *AcPIP1-3* | LOC109721095 |
| *AcPIP1-4* | LOC109727137 |
| *AcPIP2-1* | LOC109707937 |
| *AcPIP2-2* | LOC109715516 |
| *AcPIP2-3* | LOC109713634 |
| *AcPIP2-4* | LOC109708149 |
| *AcPIP2-5* | LOC109713056 |
| *AcSYP111* | LOC109724005 |
| *AcSYP112* | LOC109705538 |
| *AcSYP121* | LOC109713508 |
| *AcSYP123* | LOC109710368 |
| *AcSYP131* | LOC109715908 |
| *AcSYP21* | LOC109727640 |
| *AcSYP22* | LOC109707486 |
| *AcSYP31* | LOC109710966 |
| *AcSYP1* | LOC109713584 |
| *AcSYP42* | LOC109727303 |
| *AcNPSN12* | LOC109727331 |
| *AcSEC20* | LOC109723551 |
| *AcBET11* | LOC109723641 |
| *AcSYP51* | LOC109722298 |
| *AcSYP61* | LOC109723183 |
| *AcSYP71* | LOC109710984 |
| *AcSFT11* | LOC109727331 |
| *AcUSE11* | LOC109723551 |
| *AcSNAP29* | LOC109721766 |
| *AcSNAP30* | LOC109712973 |
| *AcSNAP33* | LOC109708842 |
| *AcVAMP711* | LOC109711499 |
| *AcVAMP712* | LOC109710881 |
| *AcVAMP721* | LOC109715685 |
| *AcVAMP724* | LOC109718244 |
| *AcVAMP725* | LOC109710494 |
| *AcVAMP727* | LOC109724556 |
| *AcYKT61* | LOC109725941 |
| *AcSEC221* | LOC109709435 |
